# The Neural Marketplace: I. General Formalism and Linear Theory

**DOI:** 10.1101/013185

**Authors:** Sarah N. Lewis, Kenneth D. Harris

## Abstract

We propose a mathematical theory for how networks of neurons in the brain self-organize into functional networks, similarly to the self-organization of supply networks in a free-market economy. The theory is inspired by recent experimental results showing how information about changes to output synapses can travel backward along axons to affect a neuron’s inputs. In neuronal development, competition for such “retroaxonal” signals determines which neurons live and which die. We suggest that in adults, an analogous form of competition occurs between neurons, to supply their targets with appropriate information in exchange for a “payment” returned to them backward along the axon. We review experimental evidence suggesting that neurotrophins may constitute such a signaling pathway in the adult brain.

We construct a mathematical model, in which a small number of “consumer” neurons receive explicit fast error signals while a larger number of “producer” neurons compete to supply them with information, guided by retroaxonal signals from the consumers and from each other. We define a loss function to measure network performance and define the “worth” of a producer to be the increase in loss that would result if that neuron were to fall silent. We show how slow retroaxonal signals can allow producers to estimate their worth, and how these estimates allow the network to perform a form of parallel search over multiple producer cells. We validate our approximations and demonstrate the proposed learning rule using simulations.

## 1 Introduction

The brain is made of billions of neurons, which together form the world’s most powerful information-processing machine. Neurons are complex devices, and are individually capable of much more than was once supposed. But what is most remarkable is their ability to organize themselves into extremely large networks that coherently guide the behavior of an animal, and constantly learn and adapt to changing circumstances.

Recordings of individual neurons show that their firing encodes diverse types of information, from simple sensory stimuli (Hubel and Wiesel, 1959) to representations of complex features such as locations in space (O’Keefe and Dostrovsky, 1971). Each neuron produces its firing pattern by building on the work of many others. This occurs with no central point of control, suggesting that local processes carried out indpendently by single neurons, cause the network to automatically organize into a coherent information processing system.

In this manuscript, we describe a mathematical model of a new process that could contribute to the self-organisation of neurons into functional networks. This is based on a proposal for a new form of communication between neurons, that we term the ‘retroaxonal hypothesis’. The hypothesis would represent a radical new process in neuronal physiology, but is supported by substantial, if circumstantial, experimental evidence. We have previously described some of this evidence (Harris, 2008). The present article contains an updated description of the hypothesis and supporting evidence, together with a mathematical formalism that describes how it could enable self-organization of learning networks, and simulations that validate our mathematical analyses.

Before stating the hypothesis, we draw a motivating analogy with another system: the global economy. In the economy, billions of individual processing units (people and firms) self-organize into productive networks. The mechanism that enables this self-organization is the transfer of money. Money flows in the opposite direction to goods and can be conceived of as a signal that indicates to a firm that its products are useful to others. By seeking to maximize their intake of money, producers are approximately maximizing their benefit to end-consumers.

In the economy, decisions are made in a decentralized manner. This occurs because firms compete not only to sell products to consumers, but also to each other. Even for firms that sell to other businesses, income can still correlate with their utility to end consumers. For example, a car manufacturer requires steel to produce the cars it sells to consumers. By competing to sell the types of steel the car manufacters require, steel firms indirectly maximize their benefit to consumers. The car manufacturer does not need to understand the steel-making process, but simply to select the steel that best suits its needs; conversely, the steel manufacturer does not need to understand all details of car design. The market thus allows individuals of limited processing power to form networks that make products far more complex than any of them could produce individually.

The aim of this article is to suggest how a similar form of self-organization could take place in the brain. We will regard neurons as analogous to ‘entrepreneurs’ who ‘sell’ spike trains to each other. Analogies between the brain and market economy have been made before (e.g. Kwee et al., 2001; von Bartheld et al., 2001; Balduzzi, 2014). The key novelty of the current theory is a hypothesized form of communication between neurons, analogous to a payment for services rendered. We hypothesize that chemical messages passing slowly backward along axons play an analogous role to money in the economy, indicating to neurons how beneficial their output spike trains are to the organism. Although rapid communication via action potentials only occurs unidirectionally forward along axons, the fact that slow chemical messages can flow in the opposite direction has been established for many decades. In the development of the nervous system, competition for such ‘retroaxonal’ signals determines which neurons live and which die (Hamburger, 1992, 1993; Oppenheim, 1991; Buss et al., 2006). In the adult nervous system, neuronal death is rare, but retroaxonal signals are still conveyed (DiStefano et al., 1992; Zweifel et al., 2005). The present theory suggests that these signals promote a different form of competition between presynaptic neurons, to supply their targets with appropriate information in exchange for a ‘payment’ returned to them backward along the axon. We argue that this form of competition – as with competition for monetary returns in the economy – promotes the self-organization of networks to allow sophisticated information processing.

The retroaxonal hypothesis does not concern rapid learning of associations beteween stimuli and responses. Rather, it concerns how the brain develops the internal representations of complex stimuli, that later allow behavior to be tuned to those complex stimuli. This form of learning does not occur within seconds and minutes, but over days, weeks, and years. We will suggest a mechanism by which neural representations that are useful for behavior are gradually selected and consolidated, at the expense of less useful representations. Once established, other neurons build on these representations, to allow representations of still more complex stimuli and thus the performance of more complex behaviors. We suggest that this gradual buildup of ever more complex representations might allow, for example, learning to understand language that is built from words, which are in turn built from simple phonemes.

## 2 The retroaxonal hypothesis

The classical form of communication between neurons is the action potential: an electrical impulse conducted rapidly along an axon, resulting in neurotransmitter release at a synapse. In mammalian neurons *in vivo*, action potentials travel in a strictly unidirectional manner from the presynaptic to postsynaptic cell.

Action potentials are, however, just one of many ways that neurons can communicate with each other. Like cells throughout the body, neurons release and receive many signaling molecules other than classical neurotransmitters, and the release of these substances can be controlled by intracellular events other than action potentials. And although classical neurotransmitters signal unidirectionally from the presynaptic to postsynaptic cell, other signals may propagate retrogradely from postsynaptic to presynaptic (Harris, 2008).

The effects of such retrograde signals need not be restricted to the synapse where they are received, but can be cell-wide, propagated not by electrical conduction but by the (much slower) mechanical transport of signaling molecules backward along the axon. Typically, the destination of such messages is the soma and nucleus, where arriving signals are integrated by complex molecular networks, and the results of the computation broadcast to the entire cell.

### 2.1 Outline of the hypothesis

This manuscript outlines a proposed role for retroaxonal signals in adult learning, that we term the *retroaxonal hypothesis* (Harris, 2008). The retroaxonal hypothesis posits that the state of a neuron’s output synapses, through the passing of retroaxonal messages, controls the stability of the same neuron’s inputs. The hypothesis has two components:

1. *Neurons with weak output synapses show unstable input synapses*. A neuron with no functional output synapses serves no benefit to an organism, and such neurons do not survive in early development. In adulthood, we suggest that neurons with weak downstream synapses do not die, but rather exhibit instability in their input synapses. This instability will cause the firing correlates of the neuron to change, until it fires in a manner that causes downstream synapses to strengthen. We assume that a neuron’s downstream synapses strengthen specifically when the neuron’s activity provides downstream cells with useful information. If a neuron with weak outputs has unstable inputs, its firing pattern will keep changing until finds information useful for downstream cells, and thus for animal behavior.
2. *Strengthening of a neuron’s downstream synapses consolidates recent changes in the same neuron’s inputs*. By consolidating changes to its input synapses that are closely followed by a strengthening of its outputs, and reverting other input changes, a neuron will keep those specific changes that lead to it conveying more useful information.

### 2.2 Candidate molecular mechanisms

The retroaxonal hypothesis represents a radical addition to the set of mechanisms usually invoked in neuronal network models. We argue for its biological plausibility by proposing a specific set of molecular mechanisms that could underlie it. We note that the hypothesis may be correct, even if it is implemented by other molecules to those specifically proposed below. Nevertheless, identification of a plausible set of candidate molecules forms a powerful ‘existence proof’ that the retroaxonal hypothesis could indeed be employed by actual neurons. The retroaxonal hypothesis requires the following:

- Information about synaptic strength, and about recent changes in synaptic strength, must be available at presynaptic terminals.
- Signals conveying this information must pass retrogradely along the axons of cells.
- These retrograde signals must cause the recipient cells to modulate the plasticity of their input synapses.
- Neurons must be able to revert recent changes to their input synapses.

We next examine these requirements, and show that each can be plausibly met.

#### 2.2.1 Retrosynaptic signals can communicate information about synaptic strength

Our hypothesis holds that changes in the strength of a neurons output synapses control the stability of the same neuron’s inputs. But how can a cell know about the strength of its outputs, when many of the determinants of synaptic strength are set by the postsynaptic neuron? Changes in synaptic strength are usually initated postsynaptically — Hebbian plasticity, for example, requires coincident presynaptic firing and postsynaptic depolarization, which is typically detected by NMDA receptors in postsynaptic spines. Nevertheless, a great many molecules are released by the postsynaptic cell during plasticity induction, that have been shown capable of signaling information about synaptic changes to the presynaptic axon terminal. These ‘retrosynaptic’ signalling molecules include lipids such as cannabinoids (Sjöström et al., 2003); small-molecule gasses such as NO (Hardingham et al., 2013); and proteins, including neurotrophins such as BDNF (Edelmann et al., 2014). In hippocampus, BDNF secretion is increased by stimulation paradigms that cause LTP, and decreased by stimulation paradigms that cause LTD (Aicardi et al., 2004). Together, the multiple molecules whose release correlates with various forms of synaptic plasticity form a ‘population code’ that keeps the presynaptic terminal informed of detailed changes in synaptic strength.

Our hypothesis requires that retrosynaptic signals inform the presynaptic neuron about both the current strength of the synapse, and its time derivative. Experimental evidence suggests that retroaxonal messengers such as neurotrophins are not only released by tonically strong synapses, but released at particularly high rates when output synapses increase in strength (Aicardi et al., 2004). Thus, the retroaxonal flow of neurotrophins could encode both the strength of an output synapse, and its temporal derivative.

#### 2.2.2 Signals can propagate retroaxonally

‘Retroaxonal’ signals are signals passing from a cell’s axon terminals, back to the soma along the axon. Retroaxonal signals play an essential role in neuronal development, where they are carried by molecules known as *neurotrophic factors*, including the *neurotrophin* family. In the developing nervous system, neurons die if they do not receive sufficient retroaxonal neurotrophic signals (Purves, 1988). Competition for retroaxonal signals has been hypothesized to select those cells that will best serve the animal in future life, through a process analogous to Darwinian evolution (Edelman, 1987; Purves, 1988). In this form of “survival of the fittest”, fitness would be measured by the number and strength of output synapses a neuron forms on appropriate targets. By this definition, the death of “unfit” neurons can only benefit the animal: neurons with no outputs will simply consume energy and resources while providing no computational benefit.

Compared to action potentials, retroaxonal signals travel slowly, and use different mechanisms. These mechanisms have been best studied for signals induced by neurotrophins, which initiate retroaxonal transport of a ‘signaling endosome’ — a small vesicle that carries the neurotrophin molecule, the activated receptor and other associated signaling molecules (Zweifel et al., 2005). The signaling endosome is conveyed back to the soma, where it influences gene expression, for example by activating the transcription factor CREB (cyclic AMP response element binding protein), which during development is critical for promoting neuronal survival (Riccio et al., 1999, 1997; Watson et al., 2001). In adults, neuronal death is rare, but retroaxonal communication through molecules such as neurotrophins continues (DiStefano et al., 1992). While the role of neurotrophins in local synaptic plasticity is well-studied (Poo, 2001), the role of retroaxonal neurotrophin signaling in adults is currently poorly studied. The retroaxonal hypothesis suggests a potential functional role for this process.

#### 2.2.3 Retroaxonal signals can influence plasticity of a neuron’s input synapses

While the best-known effects of retroaxonal signals are the control of cell survival, this is not their only function during development. Indeed, retroaxonal signals can also control the plasticity of a neuron’s input synapses. In the developing visual system of *Xenopus*, for example, injection of BDNF into the tectum causes a rapid upregulation of the strength of inputs to retinal ganglion cells (Du and Poo, 2004; Du et al., 2009). Retroaxonal effects on input synapses have also been demonstrated in cultured neurons of mammalian hippocampus (Fitzsimonds et al., 1997; Tao et al., 2000). In the developing sympathetic nervous system, signaling endosomes carrying retroaxonally-derived neurotrophic factors propoagate into the dendrites of the presynaptic cells, where they are able to directly promote or inhibit the formation of input synapses (Sharma et al., 2010).

#### 2.2.4 Changes to synaptic strengths can be transient

A key requirement of the retroaxonal hypothesis is that changes to synaptic strength are often transient, and that after plasticity, synapses will revert to their prior state if an active consolidation process does not occur. This requirement has been demonstrated by *in vitro* experiments. Long-term potentiation (LTP) evoked *in vitro* typically lasts only a few hours, before decaying and leaving synapses in their prior state; this transient form of plasticity is referred to as ‘early LTP’. Early LTP does not require protein synthesis, and relies instead on changes such as the phosphorylation and trafficking of AMPA receptors (Frey and Morris, 1997; Malinow and Malenka, 2002).

Early LTP can be consolidated into a permanent ‘late LTP’, by various cellular signals which typically require phosphorylation of the transcription factor CREB and subsequent protein synthesis (Silva et al., 1998; Barco et al., 2005). This consolidation only happens in synapses that have experienced early LTP, and thus exhibit a molecular signature known as a ‘synaptic tag’; untagged synapses are unaffected by consolidation (the exact molecular identity of the tag is still unclear; it may not correspond not to any single molecule, but to a coordinated set of changes including phosphorylation of CaM-kinase II and restructuring of the actin cytoskeleton inside a spine (Redondo and Morris, 2011; Rogerson et al., 2014)). If the consolidation signal is not received, tagged synapses revert to their prior state, while untagged synapses are again unaffected. A conclusion of these results is that synapses undergoing plasticity retain a memory of their previous state: if early LTP is not consolidated, synaptic strengths return after a few hours to the same value they had before potentiation began.

Could a retroaxonal signal trigger the consolidation of synaptic tags? A number of observations support this hypothesis. First, recent experiments show that synaptic tags can last for 5 hours or more under the correct conditions *in vitro* (Li et al., 2014): long enough for retroaxonal signals to propagate. Second, the molecules involved in the consolidation of synaptic tags overlap considerably with those activated by retroaxonal neurotrophin signals. The consolidation of early LTP into late LTP occurs because tagged synapses are able to capture ‘plasticity related products’ (PRPs) expressed on a cell-wide basis, leading to a permanent increase in synaptic strength in tagged synapses. The identity of the PRPs are also not fully established, but may include protein kinase M zeta, the scaffolding molecule Homer1a, and the neurotrophin BDNF (Redondo and Morris, 2011; Barco et al., 2005). Importantly, the key signal promoting the production of PRPs is activation of CREB, which appears to act primarily through synthesis of BDNF (Barco et al., 2005). Thus, activation of CREB by retroaxonal neurotrophin signals would be expected to promote cell-wide consolidation of synaptic tags.

#### 2.2.5 Specific candidate mechanism

The data reviewed above suggest a candidate mechanism for the retroaxonal hypothesis:

- Neurons release BDNF retrosynaptically across synapses which are strong, or which have been recently strengthened.
- The receipt of BDNF at an axon terminal leads to retroaxonal transport of signaling endosomes.
- These signaling endosomes pass on to the dendrites, where they cause consolidation of early LTP to late LTP in tagged synapses.
- Signaling endosomes arriving retroaxonally to the nucleus trigger gene expression of BDNF and other PRPs. These amplify the retroaxonal signal and allow consolidation of early to late LTP throughout the dentritic tree of the neuron.
- The signal can be passed on to the neuron’s own presynaptic partners, forming a recursive chain that ensures that upstream activity is also stable.
- A neuron with only weak outputs receives and synthesizes little BDNF, making its inputs permanently unstable. (Consistent with this idea, a recent study showed that cell-Specific knockdown of BDNF causes cells to have smaller spines, a signature of unstable synaptic inputs (English et al., 2012)).

We emphasize that this proposed mechanism remains a hypothesis, that has not been directly tested experimentally. Yet, it should at least serve to show that the phenomena we require for the marketplace scheme — though not found in current computational models of the brain — are well within the demonstrated physiological capabilities of neurons.

### 2.3 Where the buck stops: ‘end-consumers’

In the economic analogy, we think of neurons as producers of an information product which they ‘sell’ to downstream cells. But who is the final consumer? And how do these ‘consumer’ cells judge what information they need from producers, in order to usefully guide animal behavior?

In our framework, a consumer neuron is any cell that receives a *training signal* — i.e. a synaptically conveyed signal that rapidly and directly guides the plasticity of its other input synapses, in a manner that leads to the learning of appropriate behaviors. We refer to neurons that receive no direct training input as ‘producers’: in the theory, the role of these cells is to produce information required by the consumer cells.

The precise form of plasticity employed by consumer cells is not critical for the retroaxonal hypothesis, provided one condition holds: that *consumer neurons only strengthen input synapses that provide them with useful information*. In the economic analogy, one could say that the end consumers ‘buy only what they need’. Within this constraint, the retroaxonal hypothesis is agnostic about how consumers learn. Animals learn by many different mechanisms, each of which presumably requires a unique form of plasticity rules in consumer neurons. Some of these learning mechanisms are reasonably well understood (such as reinforcement learning and classical conditioning), while others are currently mysterious (such as the way that animals learn by observing the behaviors of others). These disparate learning mechanisms are believed to operate through the action of Specific neural pathways carrying training signals, for example the dopamine system in the case of reinforcement learning (Schultz et al., 1997).

Consistent with a partition of cells into ‘producers’ and ‘consumers’, training signals do target Specific neuronal populations: for example, while the basal ganglia receive extremely strong dopaminergic innvervation, the hippocampus and neocortex (particularly sensory cortex), receive far less (Bentivoglio and Morelli, 2005). We might regard the cells in basal ganglia that receive the dopaminergic training signal to be ‘consumers’, and the neocortical neurons that provide their inputs to be ‘producers’.

### 2.4 Two possible examples of retroaxonal learning

To clarify how the neural marketplace theory would operate in actual brain circuits, we now consider two Specific examples.

#### 2.4.1 Fear conditioning

Our first example is classical fear conditioning, where we suggest that retroaxonal signals could contribute to stabilizing representations of a ‘conditioned stimulus’ associated with fear memory.

In classical fear condition, pairing of an aversive unconditioned stimulus (US) with a conditioned stimulus (CS) leads to the CS evoking fear-related behaviors such as freezing. Animals can rapidly learn to fear multiple forms of CS, from simple sensory stimuli such as a previously neutral tone (auditory fear conditioning), to complex cue combinations such as those indicating particular spatial locations (contextual fear conditioniong). A considerable body of evidence suggests that fear conditioning occurs through Hebbian synaptic plasticity in the amygdala: coincident firing of a strong input signaling the US, together with initially weak synaptic inputs signaling a CS leads to strengthening of the synapses carrying the CS, after which the CS can drive a response alone (Pape and Pare, 2010). In our theory we would class the amygdalar cells exhibiting this Hebbian plasticity as ‘consumers’, with the strong input encoding the US as their training signal. Information about the CS can come from many different structures including the auditory cortex and thalamus (for auditory fear conditioning) and hippocampus (for contextual conditioning to a particular location in space). We regard cells in those areas as ‘producers’.

According to the retroaxonal hypothesis, after fear conditioning has strengthened synapses carrying the CS to the amygdala, a slow retroaxonal signal will pass to the subset of presynaptic neurons whose outputs signal the CS, ensuring that the representation of the CS in these upstream areas remains stable, while representations of other, irrelevant, features are less stable. Thus, we might expect synaptic consolidation in auditory areas when the CS is a sound, in hippocampal place cells when the CS is a location in space, and so on.

Results of contextual and auditory fear conditioning experiments are consistent with such Specific synaptic consolidation. When an animal visits a new location, place representations by hippocampal neurons form rapidly (Frank et al., 2004), but are often unstable (Kentros et al., 2004; Agnihotri et al., 2004; Kentros et al., 1998). After contextual fear conditioning, synapses from neurons representing the conditioned location onto neurons mediating fear behavior in the amygdala become strengthened (Anagnostaras et al., 2001). We suggest that this strengthening causes a retroaxonal signal to pass from the amygdala to those precise hippocampal cells encoding the conditioned location, which will stabilize their place fields and ensure that the representation of this location is retained over the long term. Consistent with this hypothesis, a wave of CREB phosphorylation (a signature of synaptic consolidation) is found in the hippocampus several hours after contextual fear conditioning (Trifilieff et al., 2006). This delayed CREB phosphorylation is what would be expected from a slow retroaxonal signal from amygdala to hippocampus. Moreover, this wave of hippocampal CREB phosphorylation is *not* seen after auditory fear conditioning (Trifilieff et al., 2006), which suggests that it follows Specifically from potentiation of inputs from the hippocampus to the amygdala.

In auditory fear conditioning, the CS is a sound. Some information about the CS arrives at the amygdala from the thalamus, Specifically the medial geniculate nucleus (MGN), and strengthening of synapses between MGN and amygdala is believed to underlie auditory fear conditioning (Pape and Pare, 2010). The MGN projects directly to the basolateral amygdala (BLA) but not vice versa. During auditory fear conditioning in rats, there is plasticity in both BLA and MGN (Maren et al., 2001). Despite the lack of direct anterograde connections from BLA to MGN, the plasticity in MGN is suppressed if BLA is silenced during training using muscimol. We suggest that retroaxonal signals from BLA to MGN, dependent on the strength of synapses from MGN onto BLA, contribute to this process.

#### 2.4.2 Reinforcement learning

The ‘law of effect’ (Thorndike, 1898, 1911) states that if an animal performs a particular action in a particular situation, and this is followed by a rewarding outcome, then the probability the action will be performed in this situation increases. It is believed that dopamine signaling is fundamental to this process, with dopamine release indicating increases in expected future reward (Schultz et al., 1997). We regard the recipients of this dopamine signal as ‘consumers’, and cells providing their inputs (but receiving no dopamine signal) as ‘producers’.

The basal ganglia are believed to be central to reinforcement learning (Ito and Doya, 2011), and are heavily innervated by dopamine (Bentivoglio and Morelli, 2005), whereas dopamine innervation of sensory cortex is weak. Although the exact mechanisms of reinforcement learning are still debated, most current hypotheses center on plasticity of the corticostriatal synapse (Costa, 2007; Wickens, 2009; Fee, 2014). It is well established that dopamine controls corticostriatal plasticity (Lovinger, 2010), and theoretical models have suggested how dopamine-gated plasticity might enable reinforcement learning to occur (Wickens, 2009; Fee, 2014; Ito and Doya, 2011; Frank, 2011). It is widely held that corticostriatal projection neurons encode sensory and contextual factors; that firing of specific striatal neurons causes production of specific behaviors; and that strengthening of corticostriatal synapses thereby increases the probability that a Specific behavior will be performed in a specific circumstance. In the present theory, we would consider the striatal cells, which receive a dopaminergic training signal, as ‘consumers’; corticostriatal projection neurons would be classed as ‘producers’. Despite their lack of a direct training signal, we hypothesize that corticostriatal neurons would receive indirect, slow reinforcement, in the form of retroaxonal signals from the striatum. According to the hypothesis, strengthening of a corticostriatal synapse would cause a retroaxonal signal to pass to the cortical neuron, stabilizing representations of those cortical neurons that encode behaviourally relevant features. Meanwhile, other cortical neurons would continue to change their input synapses until they found a behaviorally relevant variable to encode.

### 2.5 Unsupervised learning and the retroaxonal hypothesis

In section 2.4, we hypothesized that areas such as cortex and hippocampus contain populations of ‘producer’ neurons, whose plasticity is not strongly controlled by training signals such as dopamine. Instead, these producers continue to experiment with different input weights until they produce a signal that is used by the ‘consumer’ neurons.

How might producers select candidate input weights? For the retroaxonal learning scheme to work efficiently, we believe producers must use *unsupervised learning rules*. An exhaustive or random search over the very large space of possible input synaptic weights would be too slow. Instead, a producer neuron should seek representations of its inputs that are *a priori* likely to be useful. This task — finding salient features in high-dimensional data without a direct training signal — is precisely what unsupervised learning algorithms are designed to accomplish.

Unsupervised learning is frequently used in machine learning to form low-dimensional representations of complex data sets. Artificial unsupervised algorithms include principal component analysis, independent component analysis, and cluster analysis, all of which can be implemented neurally as Hebbian learning rules (Hinton and Sejnowski, 1999). Unsupervised learning has long been suggested as a key computational function of the neocortex (Marr, 1970), and several details of cortical synaptic plasticity rules appear consistent with a role in unsupervised learning (Cooper et al., 2004; Clopath et al., 2010; Yger and Harris, 2013). Implementation of synaptic plasticity rules consistent with the physiology and molecular mechanisms of cortical synaptic plasticity can allow simulated recurrent networks to form unsupervised representations of real-world stimuli such as speech sounds (Yger and Harris, 2013).

#### 2.5.1 Unsupervised learning by hippocampal place cells

Returning to the example in section 2.4.1, we note that recent data on rat hippocampal place cells suggest one way that unsupervised learning might operate *in vivo*.

When a rat is introduced into a new spatial environment, place fields representing locations in this environment appear essentially instantaneously (Hill, 1978; Wilson and McNaughton, 1993; Frank et al., 2004). These place fields are initially coarse but over a period of minutes, they grow smaller, tighter, and more reliable (Wilson and McNaughton, 1993; Frank et al., 2004).

The initial appearance of place fields appears to result from a reduction of synaptic inhibition. When a rat is introduced to a novel environment, the firing rate of putative fast-spiking interneurons drops rapidly (Wilson and McNaughton, 1993; Nitz and McNaughton, 2004; Frank et al., 2004), but returns to baseline during subsequent exploration sessions. By allowing previously sub-threshold inputs to drive spiking, this decrease in inhibition may allow the cell to fire with some degree of spatial specificity when an animal first visits a new environment, even before any synaptic plasticity has taken place. This possibility is supported by computational models, which show that a hippocampal neuron with even a randomly-weighted combination of inputs from entorhinal grid cells would show some spatial specificity (de Almeida et al., 2009), and is further supported by the observation that injecting depolarizing current reveals spatially specific firing even in hippocampal neurons that showed prior no place-related activity (Lee et al., 2012). It seems likely that refinement of place fields after their initial formation happens via Hebbian LTP of each place cell’s input synapses (Solstad et al., 2006; Franzius et al., 2007).

Thus, it appears that a coarse place field generated by random initial connectivity can form a ‘seed’ from which Hebbian plasticity sculpts a coherent place field. This unsupervised learning strategy, requiring no explicit training signal, enables ‘producer’ place cells of the hippocampus to form representations of space that are potentially useful.

Experimentally, place fields are sometimes stable from one day to the next, and sometimes not (Kentros et al., 2004). The retroaxonal hypothesis suggests that the stability of a place field corresponds to its utility to the animal: cells encoding place fields useful to downstream computations will form strong synapses onto downstream consumer cells, and in return receive a retroaxonal ‘payment’ which causes them to stabilise their input synapses. In support of this idea, most of the novel place fields formed after exposure to a novel environment disappear after one or two days; furthermore, the place fields that remain tend to be those that encode the most spatial information (Karlsson and Frank, 2008).

#### 2.5.2 Non-retroaxonal cues for producer plasticity

Retroaxonal signals could interact with other cues for producer plasticity. While the sensory cortices and hippocampus receive far less dopamine innervation than the striatum, they are strongly innervated by cholinergic fibers (Frotscher and Léránth, 1985; Eckenstein et al., 1988), which are active at times of high alertness, and lead to increased synaptic plasticity (Hasselmo, 2006). Stimuli occurring at times of high alertness are more likely to be behaviorally relevant. Thus, neuromodulators may signal to producer cells that forming a long-lasting representation of their current inputs may be useful to their downstream targets at some time in the future. Nevertheless, uniform plasticity of all neurons receiving cholinergic signals could cause older, useful representations to be lost. If retroaxonal signals dictate which cells become plastic in response to this cholinergic signal, it would protect old, useful representations, while allowing new representations to be formed.

### 2.6 How does a neuron know which changes to consolidate?

Retroaxonal signals are conveyed by physical transport of chemical signals, rather than electrical pulses, and thus are likely to take minutes or hours to travel back along a reasonably-sized axon. The slowness of this transmission raises a question: if a neuron had made multiple changes to its inputs before the arrival of a retroaxonal message signaling consequent changes to output synapses, how would the cell know which input change was responsible? At least in the case of hippocampal place fields, the question has a false premise.

When a rat is introduced to a new environment, most neurons do not form new place fields; and the few that do only form one new place field (Hill, 1978; Wilson and McNaughton, 1993; Frank et al., 2004). Thus, each cell which does form a new place field makes a single coherent set of changes to multiple input synapses, that in turn endows it with a coherent representation of a single novel variable. The changes to a single cell should therefore be consolidated or rejected all together. Because producer cells make these changes in input synapses only rarely, they will not make more than one coherent large-scale change during the time it would take a retroaxonal signal to be conveyed back to the soma.

### 2.7 Relation to other theories of network plasticity

Before describing the mathematical theory in detail, we first consider its relation to other common mathematical models of network plasticity.

#### 2.7.1 Retroaxonal learning as a complement, not an alternative

Most current theories of network plasticity concern how synaptic weights adjust rapidly to enable learning of tasks. The present theory provides not an alternative, but an addition to these theories. In the neural marketplace hypothesis, rapid plasticity occurs through conventional mechanisms, such as Hebbian unsupervised learning, supervised learning, and reinforcement learning, that have been extensively modeled already. The key novelty here is a second, slower, ‘consolidation’ process, which selectively stabilises the more useful results of those conventional learning mechanisms. The retroaxonal hypothesis is not a model of how we learn new information in minutes or hours, but rather for how skills might be built up over days, weeks, months or years.

#### 2.7.2 Fast global reinforcement versus slow, cell-specific reinforcement

Reinforcement learning guided by a single reward signal to the whole network suffers from a ‘credit assignment’ problem: if a large number of neurons or synapses make simultaneous changes, it is unclear which change was responsible for the improvement in performance. A number of solutions have been proposed (e.g. Friedrich et al., 2011; Urbanczik, 2009; Vasilaki et al., 2009; Izhikevich, 2007; Fremaux et al., 2010; Florian, 2007; O’Brien and Srinivasa, 2012), which typically involve modulation of local synaptic plasticity by a global reinforcement signal, biologically assumed to correspond to dopamine. While this global reinforcement signal is not *cell*-specific, it can be fast, i.e. *time*-specific. Thus, immediate rewards can modulate plasticity of synapses that were recently active. This is one way of partially solving the credit assignment problem.

The present theory proposes a complementary mechanism. In our theory, the producer neurons need not receive a direct, fast reinforcement signal such as dopamine. Instead, they receive indirect, slow feedback from the consumer neurons. We show that this feedback can act as a cell-specific reinforcement signal, telling each producer how important it is to the consumers. This theory may be particularly appropriate for areas such as sensory cortex, which receive little direct dopamine innervation or any other apparent training signal, but in which develop representations that appear tailored to an animals behavioral requirements (Sigala and Logothetis, 2002; Kuhl et al., 1992).

#### 2.7.3 The backpropagation algorithm

The artificial neural network algorithm that has historically seen most use in real-world applications, does involve retrograde signals. This algorithm is known as ‘error backpropagation’ or ‘backprop’ (Rumelhart et al., 1986). In the back-prop net, the firing of neurons is determined by classical anterograde transmission. However, synaptic plasticity in this network requires a training signal to flow backwards along the same connections, so that each neuron integrates a training signal from precisely those cells to which it sends axons.

Simple versions of backprop train a layered, feedforward network with a single ‘output node’ in the final layer. The output node receives, as a training signal, a target output *y(t)*, while its actual output is 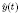. Backpropagating signals allow each upstream neuron to measure the partial derivative of 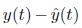 with respect to each of its input synaptic weights, and thus to do gradient descent of the mean-square loss function 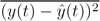. These backpropagating signals are *instructive*, meaning that they tell each cell how to change its response to its current pattern of input.

While the backprop algorithm rapidly saw success in real-world applications, it was immediately recognized that backprop was unlikely to be a model for neuronal plasticity in the brain (Crick, 1989). Although neurons are certainly capable of carrying retroaxonal signals, the retroaxonal signals found in real neurons are simply too slow. In the backprop algorithm, weight changes are based on the instantaneous correlation of presynaptic activity and the retroaxonal training signal, requiring that the training signal arrives while the input pattern is still being presented. This requires conduction in a matter of milliseconds, rather than the minutes or hours required for physical transport of retroaxonal chemical signals (Zweifel et al., 2005).

In the neural marketplace theory, retroaxonal signals are *selective*, not *instructive*: they tell each neuron *how useful it is*, rather than *how to be more useful*. The retroaxonal hypothesis requires that consumers only form strong input synapses from producers that provide useful information. Thus, to measure its utility to downstream cells, a producer neuron need only know the strength of downstream synapses, and not an instant evaluation of downstream firing patterns. Such selective retroaxonal signals need not be conveyed in milliseconds, but could take minutes, hours, or more, consistent with known biology of retroaxonal signals.

The difference between the retroaxonal hypothesis and the backpropagation algorithm can be illustrated using the economic analogy. In a market economy, consumers choose which products to buy; but it is rare for a consumer to explain to the supplier what changes in the manufacturing process would make a product more attractive. It is up to the producer to experiment, and discover what sells. Similarly, the retroaxonal hypothesis posits that a ‘producer’ cell can experiment autonomously with changes to its input synapses, and retain configurations that result in its output being ‘bought’ by downstream neurons. This contrasts with the backpropagation algorithm, where signals from downstream neurons tell each cell precisely what changes to make to its input synapses.

An additional difference between the current proposal and the backprop algorithm, is that the backprop algorithm is a gradient-based learning rule. While gradient-based optimization is guaranteed to converge to a local optimum solution, these local optima can still have poor performance compared to other solutions. The present theory is a non-gradient approach. When a producer neuron experiments with a new firing pattern, it makes large and often random changes to its inputs unrelated to the gradient of overall performance. This enables the algorithm to avoid local minima, in a manner closely related to the simulated annealing method (Kirkpatrick et al., 1983).

#### 2.7.4 Theories involving feedback connections

We have suggested that cell-specific reinforcement is conveyed to neurons by retroaxonal feedback. Could this reinforcement come instead through conventional anterograde transmission of action potentials along separate feedback pathways? There are several difficulties with this proposal. First, for some of the pathways along which we hypothesize retroaxonal signals flow — such as the corticostriatal pathway — there are no direct feedback connections. Anterograde passage of information from striatum to the sensory cortex involves an indirect, polysynaptic pathway involving multiple steps in basal ganglia, thalamus, and other cortical regions (the sensory cortex is not innervated by those thalamic regions that receive basal ganglia input). Second, even in cases where feedback connections are direct (such as from motor cortex to sensory cortex (Petreanu et al., 2012)), the feedforward and feedback projections are both highly divergent and highly convergent. Thus, if a consumer cell in motor cortex fires action potentials to signal to sensory cortex that a recent change in input provided useful information, it is entirely unclear how it would target the particular neuron in sensory cortex that made the change. Third, in cases where a role for top-down projections have been experimentally established, they appear not to be specifically involved in learning, but rather in cognitive control processes such as attention (Moore and Armstrong, 2003; Harris, 2013).

An important and popular class of theories of cortical computation suggest a specific role for top-down connections: that they subserve a ‘generative model’ predicting sensory inputs, and allowing sensory cortex to signal specifically when its inputs differ from these predictions (Mumford, 1992; Lee and Mumford, 2003; Rao and Ballard, 1999; Friston, 2008; Hinton et al., 1995, 2006); in many such theories, these feedback connections also play an essential role in sculpting plasticity in lower-level areas in an instructive manner. These theories are not inconsistent with the current proposal but rather complementary. Indeed, just as retroaxonal signals passing backward from association to sensory cortex could select neurons representing useful sensory features, retroaxonal signals passing from sensory to association areas may select association neurons providing sensory cortex with useful contextual information.

## 3 Outline of the mathematical theory

The remainder of this manuscript outlines a simplified mathematical model of the learning scheme described above. This is done at two levels of abstraction:

- At the first level of abstraction, we consider any neural network (spiking or rate-based, feedforward or recurrent) whose configuration is summarized by a matrix of synaptic weights, and whose performance can be measured by a loss function. We outline a learning scheme applicable to any such network, and define conditions that a network must satisfy for the proposed learning scheme to function.
- At the second level of abstraction, we consider a specific simple neural architecture, and give a proof that such networks satisfy the required conditions.

The proposed learning scheme centres on the idea of ‘worth’ of a cell. The worth of a cell measures the usefulness of its output, and is defined as the worsening in network performance if the cell were to die. Using the two levels of abstraction shown above, we show that:

- If neurons are able to independently estimate their worths, this allows a form of parallel search which solves the credit-assignment problem. In our proposed learning scheme, a cell is more likely to experiment with changes to its input synapses if its worth is low; and these changes will likely be transient unless they lead to an increase in the cell’s worth. We show that this promotes good performance of the network as a whole.
- In specific model networks, neurons can estimate their worths by passage of slow retroaxonal signals. Note that while the definition of worth involves the death of a cell (or the silencing of its output), cells can estimate their worth continuously without requiring actual output silencing, but by passing retroaxonal messages. Specifically, the worth of a producer cell is estimated to be large if it projects to consumer cells with strong weights; or if it projects with strong weights to other producer cells, which have large worth themselves.

The key to the learning scheme is a decentralization of decision making. Intuitively, we consider that each neuron can control the strength of its input synapses, while passively monitoring the strengths of its output synapses (which are controlled by its target cells). By changing its input synapses, a neuron changes its firing pattern. This in turn causes plasticity elsewhere in the network. If this plasticity results in the cell producing more useful information, its outputs to consumers or other producers will be strengthened, resulting in an increase in its estimated worth and indicating that these input changes resulted in improved network performance. The changes will therefore be consolidated. However if a cell already has high worth, it is unlikely that further changes to its inputs will increase its worth yet further; thus cells of high worth will undergo less frequent input plasticity. Conversely, a cell of low worth has ‘little to lose’ by experimenting with changes to its inputs, and will do so frequently.

## 4 Learning using worth in an arbitrary network

At the first level of abstraction, we consider a network of producer cells, whose free parameters are its synaptic weight matrix **W**, with elements *w_ij_* representing the strength of a synapse from cell *j* to cell *i*. The matrix **W** includes the weights from external inputs to producer cells and recurrent weights amongst produces, but not the weights from producer cells to consumer cells. We assume that consumer cells operate a learning rule to derive optimal producer to consumer weights **V**(**W**), dependent on the firing patterns of producer cells, and thus ultimately on the producer weight matrix **W**. The performance of the network depends on the output of the consumer cells, and is measured by a loss function *L*(**W, V**). Assuming that **V** is at its optimal value given **W**, the loss function can therefore be expressed as a function of the producer weights only, *L*(**W**) = *L*(**W**, **V**(**W**)). At this level of abstraction, the network could consist of either spiking or rate neurons and have a recurrent or feedforward architecture, and the loss function could be arbitrary; a specific example is given below in section 5.

### 4.1 Defining the ‘worth’ of a cell

Central to the mathematical theory is the concept of the *worth* of a neuron (or set of neurons), which we define as the amount that the loss function would increase if the neuron (or set) fell silent. In order to make a formal definition, we introduce some notation. We denote sets of cells by blackboard bold capital letters: 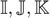, and so on. If 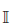 is a set of neurons, then 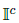 denotes the complement of this set. We write **w**_*i*_ for the vector of input weights to cell *i*, and 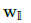 for the submatrix of input weights to a set of cells 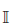. If we partition all our neurons into discrete subsets 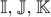, we can write the loss as 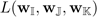, to consider how it depends on these subsets individually. We write 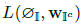 for the performance when cells 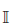 are silenced.

**Figure 1:**
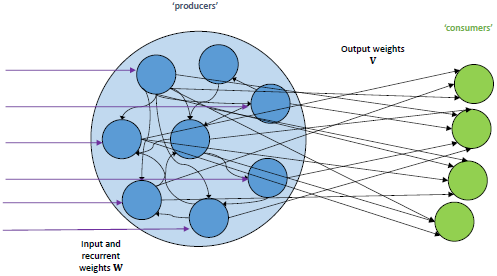
Generic network architecture. Weights onto producer neurons (from external inputs or other producers) are summarized in a matrix **W**. Weights onto consumers are summarized by a matrix **V**.

#### Definition 1 (Worth)

*The worth of 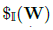 of a set of cells 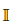 is the increase in loss function if these cells were to die (in other words, if their outputs were to fall silent):*
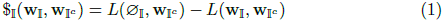

Silencing one neuron may lead to compensatory plasticity elsewhere in the network: for example if the neuron is part of an assembly of neurons with strongly correlated outputs, the silencing of its output could be mitigated by amplifying the outputs of other neurons in the assembly. We define worth to be the change in loss *after* such compensatory plasticity has occurred. For the present analysis, we assume that such compensatory changes occur only in the inputs of consumer cells **V** and not the other producer weights 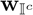.

We will show later that in specific network architectures, neurons may estimate their worth through retroaxonal signaling. For now however, we simply assume that neurons have a way to estimate their worths, and investigate the consequences of this for network learning schemes.

### 4.2 Cells should aim to increase their worth

If neurons individually change their input synapses in such a way that increases their worths, they are able to to increase the performance of the whole network. This idea is formalized as follows:

#### Theorem 1

*When a set of neurons* 
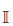
 *change their input synapses, the resulting change in network performance is equal to the change the worth of those cells*.

Intuitively, worth measures the amount that any one neuron is ‘useful’ to the broader network, and by increasing their usefulness, neurons increase overall network performance.

*Proof*. Suppose that the neurons 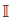 change their input weights from 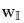 to 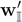.

By the definition of worth:

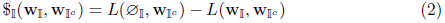

Similarly,

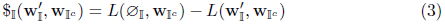

Subtracting these equations, we obtain:

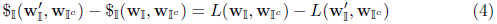

which is the desired result.

### 4.3 ‘Plasticity’ and ‘consolidation’ phases

We describe a scheme which operates by iteration of two phases: a plasticity phase, and a consolidation phase. Each phase uses measurements of worth in a different way.

#### Plasticity phase

A subset 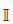 of producer neurons make experimental changes to their input synapses, each 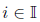 changing its input weights from 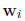 to some new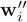. A cell is more likely to experiment with a change (‘propose a change’) if its worth is low, as making changes to these cells’ inputs is more likely to improve performance. The selection of each new 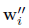 may be random, or may be guided by an unsupervised learning rule. Simultaneously, consumer neurons adjust to altered output patterns of producer cells by adapting their input weights **V** as directed by their training signals.

#### Consolidation phase

Each producer neuron forms an updated estimate of its worth, via slow retroaxonal signals. If a cell in 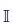 estimates that its worth has increased following the change, it retains the change, keeping input weights 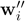; if it estimates that its worth has decreased, it is likely to revert those changes and restore old input weights **w**_*i*_. At a biological level, this ‘consolidation phase’ corresponds to the conversion of early **LTP** to late **LTP** (Redondo and Morris, 2011; Rogerson et al., 2014).

### 4.4 The Metropolis-Hastings algorithm

We formalize our proposed learning scheme as a variation of the Metropolis-Hastings algorithm (Metropolis et al., 1953; MacKay, 2003), which is at the heart of stochastic optimization schemes like simulated annealing (Kirkpatrick et al., 1983). Like our proposed learning scheme, the generic Metropolis-Hastings algorithm cycles through two steps: ‘proposing’ a new value (in our case, new synaptic weights) and then ‘accepting’ or ‘rejecting’ this change.

The Metropolis-Hastings algorithm is a way to sample from an arbitrary ‘target’ probability distribution. For a given target distribution *p*(**W**) the algorithm is as follows:

#### Algorithm 1 Generic Metropolis-Hastings

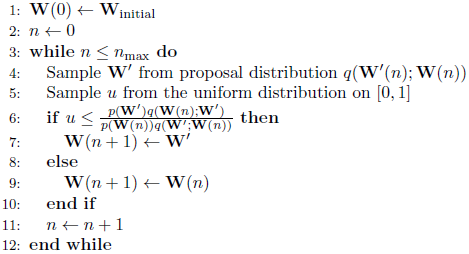

Subject to mild conditions on *q*, this defines a Markov chain on **W** which satisfies detailed balance for *p*, and as 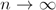, as the distribution of **W**(*n*) tends to *p*. There is no guarantee as to the distribution of **W**′.

#### Target distribution for ‘learning’

The aim of learning is to adjust synapses so that the loss *L*(**W**) is lowered. Therefore, we choose as our target distribution 
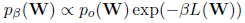
where *β* is some positive number and *p*_0_ encodes constraints on **W** (for example, that the output weights of excitatory cells must be positive). Sampling from this distribution will stochastically explore different values of **W**, preferring values that make *L* small.

The parameter *β* is known as ‘inverse temperature’. When *β* is large, the algorithm becomes a greedy optimisation of *L*, whereas with *β* = 0 it will simply sample from *p*_o_, ignoring *L*. (The process of running the Metropolis-Hastings algorithm while progressively raising *β*, known as simulated annealing, is a common method of searching for global optima of analytically intractable objective functions.)

With this choice of target distribution, the ‘acceptance probability’, i.e. the chance of executing line (7) in algorithm (1) becomes 
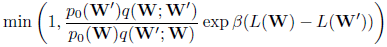

If *q* satisfies detailed balance for *p*_0_, i.e. 
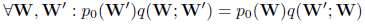
then the acceptance probability is simply 
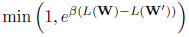
and algorithm (1) becomes

##### Algorithm 2 Simple Metropolis-Hastings

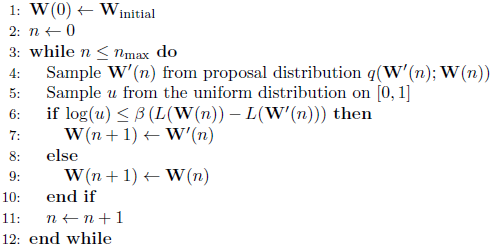

One can imagine a way the brain could employ Algorithm (2). During learning, an animal experiments with a change to the weights of the synapses of multiple neurons, proposing new weights **W**′ drawn from 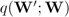. Then by attempting to perform a behavioral task, the animal determines whether the loss function *L* has increased or decreased. If loss has decreased, all the changes are retained, but if loss has increased the weights are likely to revert to **W**. One could imagine a ‘consolidation signal’ being encoded by global dopamine release, occurring with probability 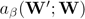. Such a scheme would be compatible with numerous aspects of known neurophysiology: the ‘synaptic tagging’ literature shows that changes to synaptic weights are usually transient unless consolidated by a second signal; and dopamine is one of several signals able to cause this consolidation (Huang and Kandel, 1995; Sajikumar and Frey, 2004; Kentros et al., 2004).

#### Chimeric acceptance

While the scheme just described would theoretically, eventually, converge to the desired probability distribution *p_β_*(**W**), it would be severely slowed by a credit assignment problem. When changes are proposed to multiple cells, these are accepted or rejected wholesale, even if some changes were for the better and some for the worse. We suggest instead that, after changes are proposed to a set of cells 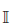, they may be consolidated by a *proper subset* of 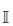. This is no longer strictly a Metropolis-Hastings algorithm because **W**(*n* + 1) may be neither **W**(*n*) nor the proposed 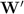, but a chimera.

If chimeric acceptance is allowed, how should each cell decide whether or not to consolidate a change it has made? From Theorem (1) we know that a useful change is likely to be one which increased the worth of a cell. Therefore, cells should generally accept changes that increase their worth and reject changes that decrease their worth. Each producer neuron uses an estimate of its worth as a cell-specific reinforcement signal.

Biologically, the worth signals could not be carried by a global neuromodulatory system such as dopamine, since worth signals are different for each neuron. We propose that this scheme can operate in brain areas such as sensory cortex, that receive little dopaminergic innervation, with worth instead conveyed to each neuron individually by retroaxonal signals.

#### Choice of proposal distribution *q*

The precise form of *q* does not affect whether the algorithm obtains the correct stationary distribution, but a good choice of *q* is important for rapid convergence. Specifically, convergence will be slow if either the proposed steps 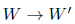 are too small, or if the proposals are too often rejected. In our case, we need to propose changes to synapses that are *a priori* likely to improve performance (and, therefore, likely to be accepted). We suggest two ways to promote this:

- Propose changes more readily to cells that have low worth.
- Use an unsupervised learning rule to propose new weights that are likely to be useful.

### 4.5 Proposed algorithm: Parallel Metropolis-Hastings (PMH)

We call our proposed learning scheme **‘PMH’** (short for ‘parallel Metropolis-Hastings’). At each step of the **PMH** algorithm the synaptic weight matrix **W**(*n*) is updated to a new value **W**(*n* + 1), via three stages:

1. each neuron decides whether to propose a change to its input weights;
2. each neuron that is proposing a change selects new input weights 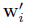;
3. each neuron that proposed a change decides whether to consolidate the change (set 
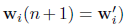
 or reject the change (set **w**_*i*_(*n* + 1) = **w**_*i*_(*n*)).

The algorithm is given explicitly in pseudocode Algorithm (3).

#### Convergence of PMH

It is not obvious that **PMH** satisfies detailed balance for *p_β_*. Indeed, it fails to do so unless certain conditions are met. In section (4.5.1) we define sufficient conditions. In section (5), we analytically determine some situations in which these conditions are approximately met, and in section (6) we perform simulations to show that the scheme works in practice. Note that the proposal distributions *q_i_* in Algorithm (3) do not correspond exactly to marginals of the proposal distribution *q* in Algorithms (1,2): for the two to correspond, we would have to include the chance of proposing a change, 
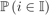
, as a factor of *q_i_*.

#### Parameters *α*, *k*

We use the parameter *α*∈[0,*β*] to dictate how strongly 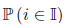 depends on the worth 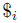, and use the parameter *κ* to control the total size of 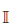. In practice, the values of *α* and *κ* may be chosen dynamically, to ensure that on each iteration 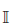 is small enough for the algorithm to converge, but large enough to allow rapid learning.

#### Why not set *α* = 0?

Why do we make the chance of proposing a change higher for cells of lower worth, when the Metropolis-Hastings algorithm does not require such discrimination?

- The Metropolis-Hastings algorithm makes no guarantee as to the distribution of proposed values 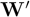, merely the stationary distribution of *accepted* values **W**(*n*). In an implementation of the scheme, proposed new weights *are* actually used for a short time, even if they then revert. A random change to a cell of high worth is likely to hinder network performance. Thus, we are likely to incur a transient penalty while evaluating changes to cells of high worth.

##### Algorithm 3 Parallel Metropolis-Hastings (PMH)

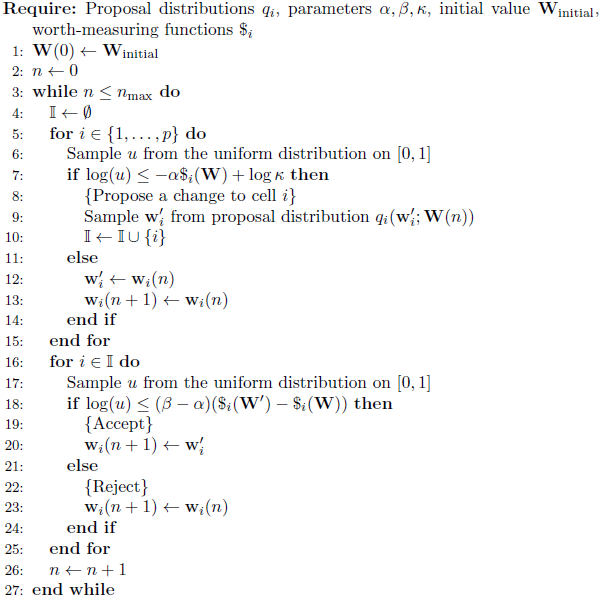

- There is an ‘opportunity cost’ to experimenting with a cell that is unlikely to improve. We will see in section (4.5.1) that our algorithm only allows us to experiment with changes to a limited number of cells at any one time. This limited set should include cells that are likely to improve.
- In our implementations of PMH (section 6), worths are estimated rather than measured exactly, and it takes some time to arrive at a good estimate. A cell which has not changed for a long time knows its worth 
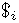
 fairly accurately. When it proposes new weights, it has only a limited time to evaluate its new worth 
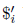
 before deciding whether to consolidate the changes. By raising *α* towards *β* we transfer emphasis from the accept/reject decision (based on a hasty estimate of 
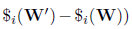
 to the proposal/no-proposal decision (based on a more accurate estimate of 
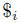
).

#### 4.5.1 Convergence of PMH

We now define sufficient conditions for the Markov process defined by PMH to have 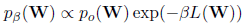 as a stationary distribution.

One condition we need is that the set of cells proposing changes at each iteration is **additive**, which we now define.

##### Definition 2 (Additive set)

*A set of neurons* 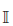 *is an* **additive set** *if for all*
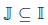 *and for all possible* 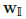, *the worth of* 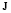 *is equal to the summed worths of the neurons within it:* 

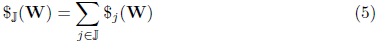

##### Definition 3 (Plus-one-additive)

*The set* 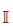 *is* **plus-one-additive** *if, for every cell i, the set* 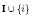 *is additive.*

For our proposed learning scheme to function, we do not require that worth is *globally* additive, i.e. that 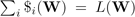; indeed, that condition is not generally met. In section 5 we will show that for a simple network, any sufficiently small set of neurons is approximately additive. In allowing each cell to decide independently whether to propose a change, we cannot strictly enforce a limit on the number that propose changes. However, we will choose small *κ* or large *α* so that in practice, the number of cells proposing changes at once is always sufficiently small.

Next we will show that additivity implies independence: if a set 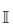 is additive, then 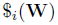 is independent of 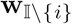. The worth of cells in an additive set 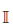 may still depend on input weights to cells not in 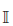: an obvious example is when cells 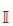 receive inputs from cells in 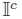.

Intuitively, additivity or independence of worths is a necessary condition for ‘credit assignment’ to make sense. If the worth of one cell 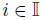 depends on the configuration of other cells in 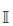, then it does not make sense to assign ‘credit’ or ‘blame’ to individual cells.

##### Theorem 2 (Additivity implies independent worths)

*Suppose that* 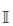 *is an additive set and* 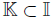. *Then the worth of set* 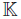 *does not depend on the weights of the other neurons* 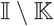.

##### Proof

Assume 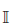 is an additive set and write 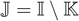.

We first show that 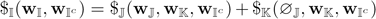. Indeed, 

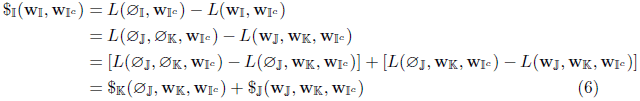

Because 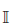 is an additive set, we also have from the definition (2) that 

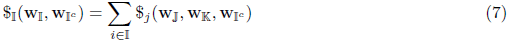

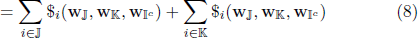

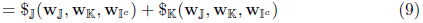

and comparing (6) to (9) we have that

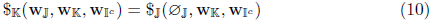

The right hand side of (10) is independent of 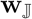, so the worth of the subset 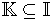 does not depend on the weights of the other neurons in 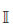.

We are now ready to state sufficient conditions under which PMH satisfies detailed balance for *p_β_*.

##### Theorem 3 (PMH satisfies detailed balance for *p_β_*)

*The transition probabilities*
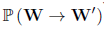 *of the PMH algorithm satisfy detailed balance for p_β_, i.e.* 

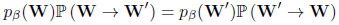

*subject to the following conditions*:

1. *The set of cells proposing changes is always plus-one-additive*.
2. *No one-way streets*: 
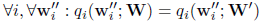
3. *Neighbours make the same proposals: whenever* 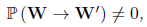 *then* 
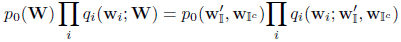
4. *Whenever* 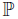 (*propose changes to 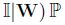* (*propose changes to 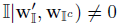 then 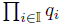 satisfies detailed balance for p_0_:* 

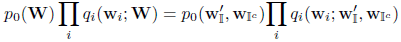

In practice, each of these conditions will only hold approximately. We therefore need to verify that the algorithm works in practice, which we do in section (6).

##### Proof

We need to show that 

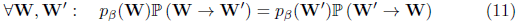

When any of 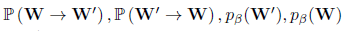 is zero, this is satisfied trivially, so let us now assume they are all non-zero.

We will write 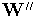 for proposed weights. 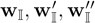, etc. denote submatrices of 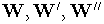 corresponding to cells 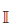. Thus, in the transition 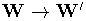, we have 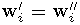 if cell *i* accepted a change and 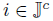 if cell *i* either did not propose a change, or proposed and rejected a change. Define

- 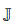 the set of cells which propose and accept changes
- 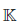 the set of cells which propose and reject changes
- 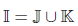 the set of cells which propose changes
- 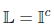 the set of cells which do not propose changes

Recall the PMH algorithm: from weights **W** one iteration is as follows

- each cell *i* decides independently whether to propose a change, with probability 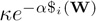.
- if cell *i* decides to propose a change, it proposes new weights 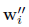 with probability 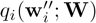.
- each cell *i* that has proposed a change decides independently whether to accept it, with probability 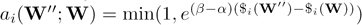.

We define 

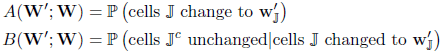

so that 

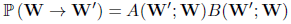

Since we assume that 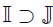 is additive and 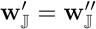, we have 

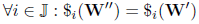

and therefore 

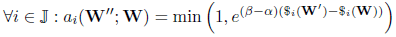

This means we can simply write: 

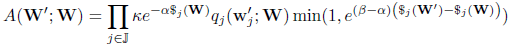

Therefore 

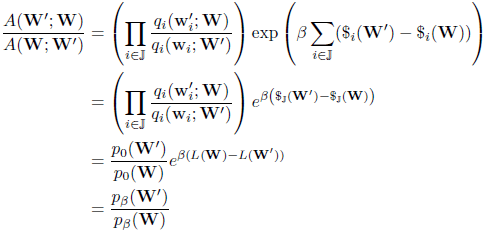

Meanwhile, *B* is the chance that each cell 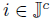 either did not propose a change, or proposed but rejected a change. Accordingly we write 

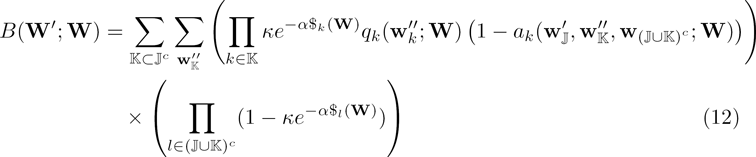

The first product 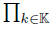… is contributed by cells that propose but reject a change, and the second product 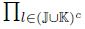… is contributed by cells that do not propose a change.

We now show that 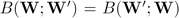. By our assumption that 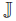 is plus-one-additive, we have that 

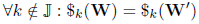

and therefore the terms 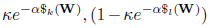 are identical for 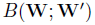 and 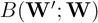. By assumption (3), the terms *q_k_*(…) are also identical for *B*(**W**';**W**) and *B*(**W**;**W**'). Finally, we need that 

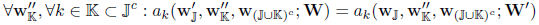

By definition 

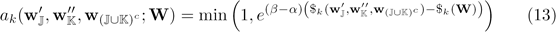

So we require that 

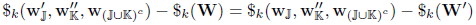

Plus-one-additivity of 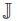 implies that 

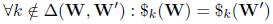

and plus-one additivity of 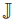 implies that 

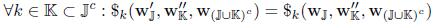

hence (13) is satisfied, and we have shown that 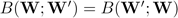. Therefore 

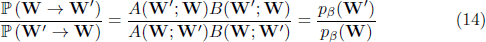

## 5 Direct worth with a linear consumer

In the previous section, we described a general learning model, PMH, in which cells use measurements of their worth to guide the plasticity of their inputs. PMH requires that cells are able to measure their worths, and that every sufficiently small set of cells is additive. We now consider a more specific model network and show that these conditions are approximately met.

We consider a model in which a pool of ‘producer’ neurons learn using PMH, while their outputs are used by a single ‘end consumer’. (The extension to multiple end-consumers is straightforward but for simplicity of notation, we consider only a single consumer). The end-consumer does not use PMH, but learns its input weights by a standard supervised learning rule, to be described below. Intuitively, we expect that producers can infer that they have high worth if their outputs are heavily used by the consumer. In this section, we validate that intuition.

In this section we discuss *direct worth:* the change in performance that would result if a producer’s output became hidden from the consumer, but not from other producers. By *indirect worth*, we mean the effect that silencing a producer would have including via its connections to other producers. Direct and indirect worth will be generally be different in recurrent networks, but equal in a feedforward architecture (Figure 2). We now show that a neuron may estimate its direct worth through the strength of its output synapses onto producer cells. A future manuscript will discuss the case of indirect worth in recurrent networks, in which case worth estimates require recursive message passing through recurrent networks.

**Figure 2:**
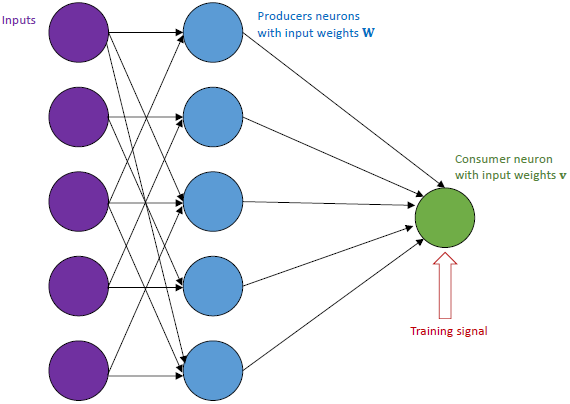
Feedforward architecture with a linear consumer. We consider a simple model in which non-linear producer neurons use PMH to learn their input weights **W**, while a single linear consumer neuron learns to approximate a target using supervised learning.

We will show how, for this model:

- strong synapses onto the consumer predict high direct worth;
- direct worths are approximately additive for small sets of cells.

### 5.1 A model consumer doing ridge regression

The consumer neuron can draw on outputs of *p* producer cells with time-varying outputs 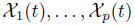, and is required to approximate a target output 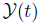 as a weighted sum 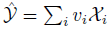, by choosing appropriate weights **v**. We define the penalized loss function *L* as 
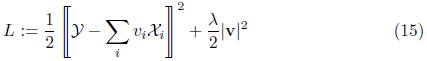

Here,

- 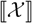 means the root mean square of *X*(*t*)
- |v|^2^ 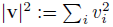, so |v| is the ordinary 2-norm.

This loss function *L* is familiar from ridge regression or penalized least squares, and the optimal **v** can be reached neurally using a delta rule with weight decay.

In general, the activity of the producer neurons 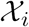, and the target function 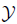 will be continuously time-varying random variables. In practice, we will train the network and evaluate its performance on finite datasets of say *N* samples. We write our *N* samples of 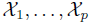 in a matrix 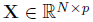 whose i^th^ column 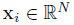 represents the output from producer cell *i*. We write the corresponding target values of 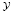 in a vector 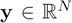, and write the output of the consumer neuron as 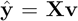. We assume that the data set is large enough to make our loss function *L* equivalent to the finite-sample loss function 

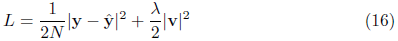

A loss function similar to equation (16) also occurs in Bayesian linear regression, where a penalty term 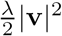 arises due to a Gaussian prior on the weight vector **v**. In the present case however, this penalty should be interpreted not as a Bayesian prior, but rather as a physical cost of synapses. The critical difference is the relative scaling of the two components of the loss function. Indeed, as *N* grows, the term 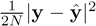 remains the same order of magnitude; thus, the penalty term 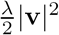 retains its importance even as the size of the data set grows. Such behavior is produced by a delta rule with fixed weight decay; but it is not the behavior that would be produced by Bayesian linear regression with a Gaussian prior on **v**, in which case the penalty becomes relatively less important as more data is acquired. Biologically, this form of weight penalty scaling might result from a physical (e.g. space or energy) cost of maintaining large synapses.

### 5.2 Ridge regression formulae

We start by recalling some formulae in the theory of ridge regression. For more details see for example (Shawe-Taylor and Cristianini, 2004; Hastie et al., 2009; Anderson, 1958)

**Optimal weights v.** At the optimal **v**, 

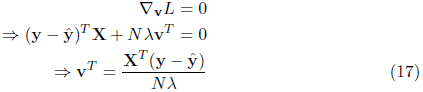

In the limit 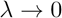, this equation implies that the residual error must be orthogonal to the activity of each producer neuron (or the weights would be infinite).

By substituting the definition 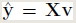 into (17), we obtain a formula for optimal weights **v** from the data **X**; **y**: 

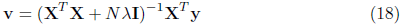

**Loss at the optimal v.** By substituting (17) into (16), we can show that the loss at the optimal value of **v** is proportional to the mean product of the training signal with the residual error: 

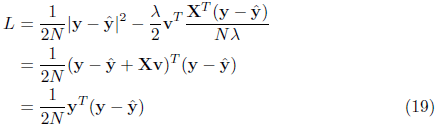

**The kernel matrix K,** the hat matrix **H**, and the residual matrix **R** We define the symmetric matrices 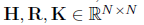 as: 

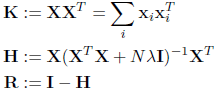

The kernel matrix **K** — which contains to the inner product of all pairs of data points — is familiar from the theory of support vector machines and other learning algorithms (Shawe-Taylor and Cristianini, 2004). **H** is called the ‘hat matrix” because it maps the target signal **y** to the consumer neuron’s actual output ‘y hat’: when the weights **v** take their optimal value, 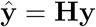. Similarly, the matrix **R** maps **y** to the residual: 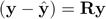.

Note that **H** and **R** depend only on **X** and not on **y**; they therefore summarize how well a given pattern of producer cell activity **X** prepares the consumer cell to learn any target pattern **y**. Specifically, **R** can be considered as a quadratic form that predicts the loss function for any given target: from (19) we have that at the optimal **v**, 

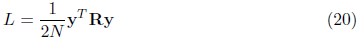

In the above equation, both *L* and **R** should be understood as having an implicit dependence on **X**.

We now derive a formula for **R** in terms of **K**. First, recall the Woodbury matrix identity, which holds that for any matrices **A**, **U**, **V** and **C:** 

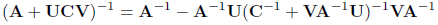

On setting 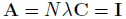 and 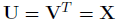 we obtain 

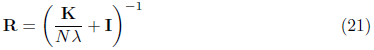

### 5.3 Direct worth is predicted by output weight to the consumer

We now show that the direct worth of a producer neuron is approximately proportional to the square of its output weight. We then consider in more detail the conditions under which the approximation holds, and argue that similar conditions apply to cortical networks *in vivo*. We first formalize the definition of direct worth:

#### Definition 4 (Direct worth)

*The direct worth Ω_i_ of producer cell i is the increase in L that results when its output weight to the consumer, v_i_, is clamped to zero, and the consumer’s other input weights v_j_ are then relearned*: 

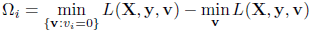

We consider the worth of producer cells as a function of their firing patterns **X**; we assume that the optimal values of the consumer’s input weights **v** are learned given **X**. If we silence the firing of cell *i* (i.e. set **x**_*i*_ = 0), without affecting the output of other producer cells, then the weight penalty in (15) means the consumer will set *v_i_* = 0. Thus, Ω_*i*_ could equivalently be defined as 

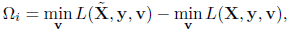

where 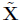 is the matrix **X** with the column **x**_*i*_ replaced by all zeroes.

We now show that the direct worth of a single cell is approximately proportional to the square of its output weight: 

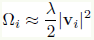

#### **Theorem 4** (Output weight predicts direct worth)

*The direct worth Ω_i_ of neuron i is given by* 

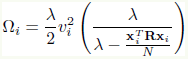

**Corollary**. If 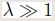 and 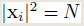 then 

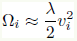

This approximation will always be an underestimate of the producer neuron’s true worth, but the estimate will be accurate when 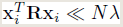 (see section 5.5 for an intuitive understanding of this condition).

#### Proof

Recall from (20) that 

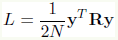

and therefore, when **X** changes in any way (and then **v** is updated), the change in loss is 

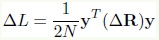

We write **R** for the residual matrix when the battery of producer cells includes cell *i* and **R**′ = **R** + **ΔR** for the residual matrix after that cell is removed. Recall that 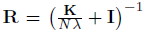, where 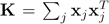, is the kernel matrix. We can therefore use the Woodbury matrix identity to compute **ΔR**. The identity is: 

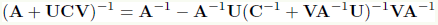

We use this with 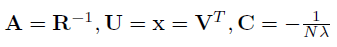 to obtain 

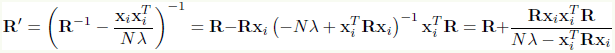

So that 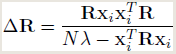

As 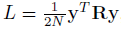, the change in *L* from removing cell *i* is 

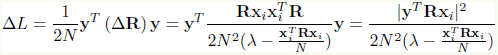

Recall also that the optimal output weight of cell *i* (before it is silenced) is 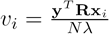. Thus, 

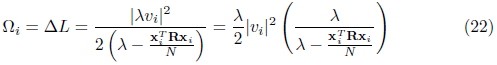

For the corollary, recall that **R** is a contraction so 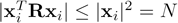; hence the factor in the second set of brackets is at most 1.

### 5.4 Direct worth is approximately additive

We now show that for small sets of cells, direct worth is approximately additive.

#### **Theorem 5** (Direct worth is approximately additive)

*Let 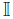be a set of producer cells, with 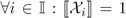. Write 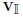 for the vector of output weights of all cells in the set 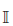, so 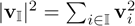. Then the direct worth 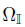 of set 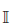 satisfies* 

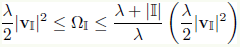

**Corollary**. If 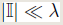 and 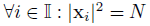 then 

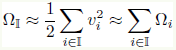

Thus, direct worth is approximately additive for sufficiently small sets of cells.

Again note that the approximation 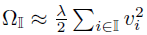 underestimates the worth of a population.

#### Proof

Again, we will use 

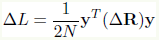

Split the producer cells into two disjoint sets 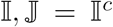. Write 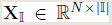 for the matrix of data from cells 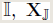 for the matrix of data from cells 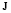. Correspondingly define 

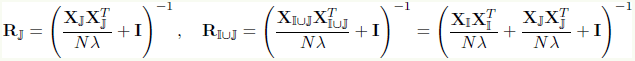

so 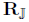 is the residual matrix when only cells 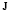 are available (see 21). Recall the Woodbury matrix identity: 

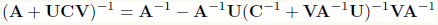

Using the Woodbury identity with 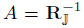 and 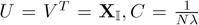, we have 

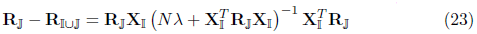

It will prove convenient to define 

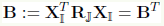
and rewrite (23) as 

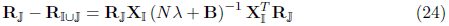

Therefore, the change in loss is 

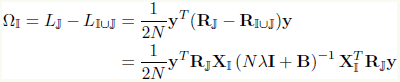

By definition the direct worth is 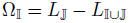 so this means that 

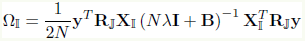

We wish to compare the direct worth to the weights of cells 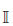. From (17) we have 

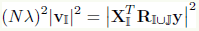

and using (23) we have 

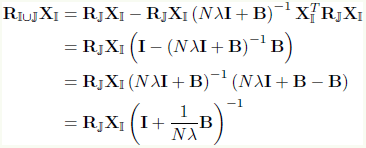

Therefore, 

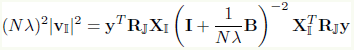

We can therefore regard 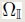 and 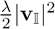 as the result of applying two different quadratic forms, 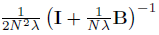 and 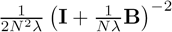, to the same vector 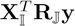. We now show that **B** has small eigenvectors so that 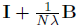 is close to identity, and thus 

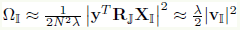

Both quadratic forms share eigenvectors with **B**, so we proceed by decomposing 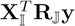 into these eigenvectors. Write *μ_i_* for the eigenvalues of **B** and **u**_*i*_ for corresponding unit eigenvectors. **B** is real and symmetric, so it has a full basis of orthogonal eigenvectors. We write 

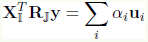

and note that 

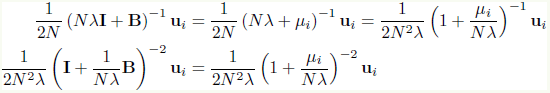

Write *μ_i_* for the eigenvalues of **B**, **u**_*i*_ for corresponding eigenvectors, and write 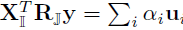. Then 

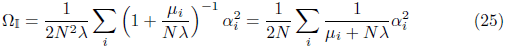

and 

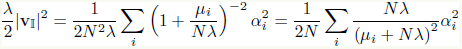

Subtracting one from the other, we obtain that the error in approximating the direct worth 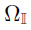 by 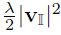 is 

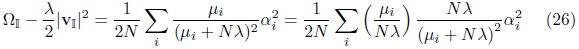

**B** is real, symmetric, and positive semidefinite, so that each *μ_i_* is real and non-negative. From (25,26) we can therefore derive the bounds 

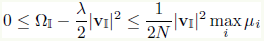

Now we show that max_*i*_ *μ_i_* is small, using Gershgorin’s theorem. This theorem says that the eigenvalues *μ_i_* of **B** must lie within ‘Gershgorin discs’ 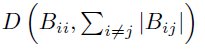 which have centers **B***_ii_* and radiuses 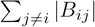.

Write 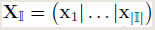 so that 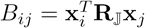. The residual matrix **R** is a contraction, so we have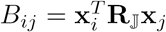. Assume that each 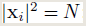; it follows that 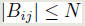. Therefore, **B** has eigenvalues inside 

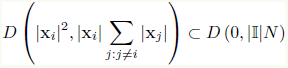

Since **B** is also positive semi-definite, it follows that each *μ_i_* satisfies 

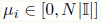

so that 

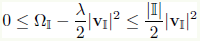

Following similar reasoning we can also bound the approximation error 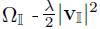 as a proportion of the exact value of 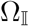: 

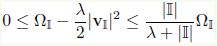

The upper bound in theorem 5 is tight. Suppose we have *p* identical cells each outputting 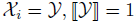. Then the weights *v_i_* satisfy 

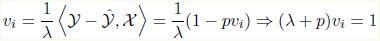

and the loss is 

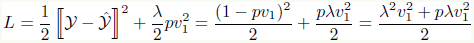

If all cells died, then the loss would be 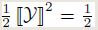. Thus, the direct worth of the set {1,…, *p*} is 

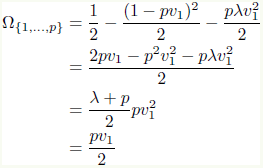

This is equal to the upper bound in theorem (5): 

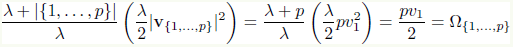

### 5.5 When do the approximations hold?

We have shown that the conditions required for the parallel Metropolis-Hastings algorithm (additivity and retroaxonal estimability of worths) can approximately hold for a linear consumer neuron. It is instructive to consider further the conditions under which these approximations hold. Loosely speaking, the answer is that the approximations hold when coding by producer neurons is highly redundant.

Without loss of generality, we assume that cell *i* has unit mean-square firing rate, i.e. 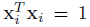. From theorem 4 we see that the approximation 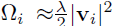 holds when 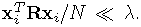. This will always be the case if 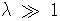; it can also hold for smaller λ, provided 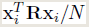 is sufficiently small. Both conditions correspond to situations of highly redundant coding. As we show below, when 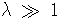, good consumer performance can only be achieved when there are large numbers of redundant producer cells encoding information that predicts **y**. Furthermore, we see from (20) that the condition that 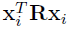 be small can be understood as a requirement that if **x**_*i*_ were presented as a target, it would be well predicted by the whole population of producer neurons. In other words, the retroaxonal estimate of worth is accurate for producer neurons that are part of a redundant assembly. We now make these ideas precise, using a singular value decomposition of the matrix **X**.

#### Singular value decomposition of X

We can gain insight into the above arguments by performing a singular value decomposition of the matrix **X**. We will see that the loss function is small when **y** is aligned with directions of high activity in **X**.

The singular value decomposition (Strang, 1993) factorizes any matrix 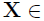
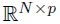 into the form 

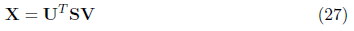

where 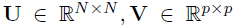 are orthonormal matrices, and 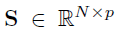 is diagonal. We denote the diagonal elements of **S** (which are called the *singular values* of **X**) as *s*_1_, *s*_2_, …, *s_N_* (padding with 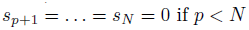).

The rows **u**_1_, **u**_2_, … of **U** — which constitute an orthonormal basis for 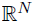will be the principal components of the data matrix **X** if the mean of each x_*i*_ is 0, and the sample variances in these principal component directions will be 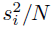.

By writing the hat matrix **H** in terms of **U**, **V**, **S** we see how to regard the map 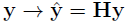geometrically. One can show that 

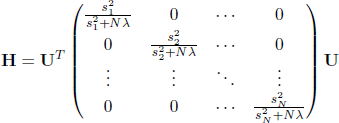

and 

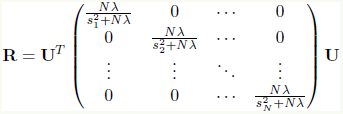

**R** is a contraction which drastically shrinks those dimensions **u**_*i*_ with 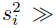
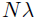, but leaves dimensions with 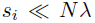 unchanged. Because the principal components **u**_*i*_ form an orthonormal basis, we can express any target vector as a weighted sum 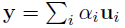. Recalling from (20) that 

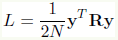
we see that 

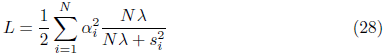

Thus, the performance of the network is good when large coefficients *α_i_* coincide with large values of 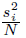. In other words, the network is able to learn efficiently if most of the length of target vector **y** is in directions of high activity in the producer cell activity matrix **X**.

#### How many producers are needed for good performance?

A sufficient condition for our approximations to hold is that 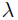 be large. We now show that good performance can only be obtained in this case through highly redundant coding.

First, suppose a set of *p* producer neurons all encode exact copies of the output target **y**, which has a mean-squared firing rate of 1. Perfect target prediction would be obtained from any weight vector **v** with 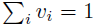. Amongst such vectors, the one which will incur the smallest weight penalty would have equal weights from all producer neurons. Note that this occurs only for an *L*_2_ weight penalty; an *L*_1_ penalty would penalize all such vectors equally. Informally, one could say the *L*_2_ norm has a ‘democratic’ property that ensures it weights all producers equally: if 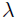 is large, the squared weight penalty dissuades the consumer from forming a strong input synapse from any single producer. This feature — that no one neuron can monopolize the activity of another — is critical for the success of the neural marketplace algorithm. It is also an ubiquitous feature of mammalian brain circuits (with the exception of a small number of ‘detonator synapses’, such as are found in the subcortical auditory system).

In practice, the optimal weights will not sum to 1, and predictions will not be perfect. To compute the actual weights, note that the matrix **X** has an singular value equal to 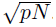. It is easy to show using (18) that the weight from each neuron is 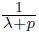, and we see from (28) that the total loss is 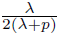. Note that loss descreases with the number of producer neurons *p*, but not with the number of training samples *N*. Thus, good performance requires that *p* ≫ λ; in other words, for large 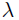, we must have redundant coding.

The aim of retroaxonal learning is to adjust the firing of producer neurons to the needs of consumers. The situation just considered — where all producers perfectly encode the target — could be considered the endpoint of this process, after which 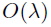 neurons are required. How many producer neurons are required for less optimal producer coding? To analyze this situation, suppose that the target and activity of all producer neurons are random combinations of the same *n* uncorrelated signals, where 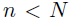, and that each has a mean-square firing rate of 1. We can think of the **x**_*i*_ (or rather, their correlations with the *n* signals) as being uniformly distributed on a unit sphere in 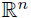. For large enough *p*, the non-zero eigenvalues of **X^T^ X** will be close to 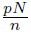 ^[1]^ Thus, the loss is approximately 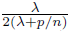, and we see that performance is good when 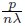 is large, i.e. when there are 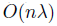 producer neurons. A retroaxonal learning rule, that selects cells highly correlated with the target, could shrink the required number of producer neurons by a factor of *n*.

#### Effects of firing rate on worth and the need for rate homeostasis

Until now, we have assumed that each producer neuron fires with a mean-square firing rate of 1; that is, 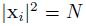. We now consider the effect of scaling a neuron’s firing rate on its worth.

As before, we assume that the consumer is able to find optimal **v** for given **X**, **y**, λ. We fix **y** and λ, and consider *L* simply as a function of **X**, using the shorthand ‘*L*(**X**)’ to mean ‘min_v_ *L*(**X**, **v**, **y**, λ). It is simple to show that for any *a* > 1, *L*(*a***X**)≤ *L*(**X**). Indeed, take **v**^*^ = argmin *L*(**X**, **v**, **y**, λ). Then for any *a* < 1 we have

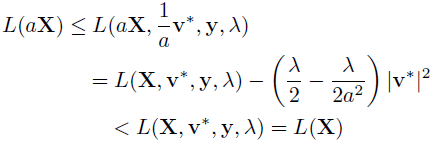

This means that a cell may increase its worth to the consumer simply by firing at a faster firing rate; intuitively one might understand this as being because a higher firing rate allows the same effect on the consumer to be achieved with a lower output weight and thus a lower penalty. To ensure that worth measures the quality, rather than quantity, of a neuron’s spikes therefore requires that a different mechanism exists to keep all cells firing in a reasonable range of rates. Such mechanisms of rate homeostasis do indeed exist in the cortex (Xue et al., 2014; Turrigiano, 2008).

## 6 Simulations

### 6.1 Direct worth

We now demonstrate the operation of the neural marketplace rule, and check the validity of our approximations in simulation. First, we check the approximation for direct worth 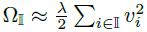using a simulation.

In this example, the output **X** of the producer neurons is generated by a random recurrent rate-based network with sigmoid nodes (Figure 3). The network is driven by a sinusoidal stimulus 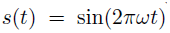, while a single ‘consumer’ node has target output oscillating at twice the frequency: 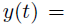
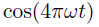. Solution of the problem is therefore only possible due to the nonlinear nature of producer neuron’s activation function (since 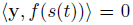 for any linear *f*). The network contains *p* = 500 neurons, excluding the single stimulus node. 20% of the neurons are inhibitory and the neurons are sparsely and randomly connected.

**Figure 3:**
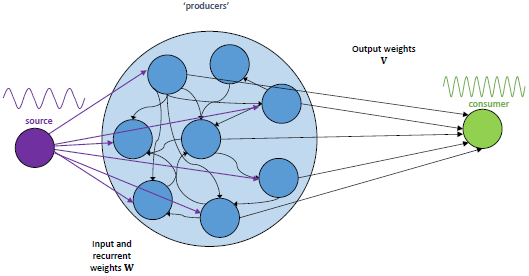
Frequency-doubling task. A single input following a sine-wave input is presented to a random recurrent network of sigmoidal units (20% inhibitory, 80% excitatory). The target output is a sine wave of twice the frequency.

A single consumer neuron attempts to approximate the target **y** as a linear combination of the producers’ outputs: it outputs 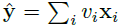 choosing **v** to minimise the loss *L* defined in (15), namely 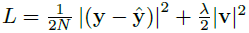.

To check the direct worth of a set of cells 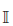 we do the following:

- Run the complete network on the stimulus for *N* timesteps and record the result in data matrix 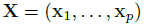.
- Find weights **v** which minimise *L* and record this *L*.
- Find weights 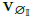 which minimise 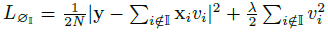 and record the corresponding 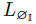.
- Record the direct worth 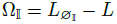.

Figure 4 shows the result of the simulations. When 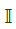 is a single cell, the results match the estimate 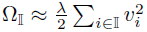 very closely. For larger sets, the approximation holds, but with quality decreasing as the size of the set 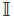 increases, showing substantial deviation when 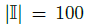 (one fifth of the total population size). The predicted worth is always an underestimate (in accordance with theorem 5).

**Figure 4:**
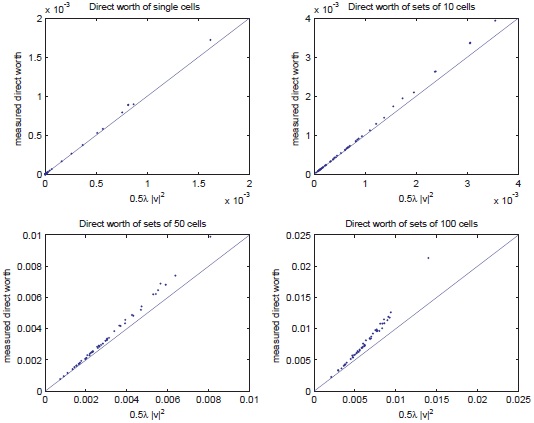
Direct worth of producer cells. The direct worth of an individual producer is very close to 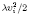. The direct worth of a set 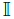 of cells is approximated by 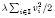, but less well.

### 6.2 PMH with unsupervised learning

We next evaluate the parallel Metropolis-Hastings scheme. Specifically, we test it on a form of ‘cocktail party problem’ (figure 5). The inputs that producers will see stimuli are linear combinations 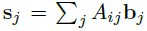 of independent source signals **b**_1_, **b**_2_,… with random weights *A_ij_*. In this simulation there are 10 consumers, and the target outputs for the consumers depend only on a small subset the source signals, with the others being distractors. This is a ‘cocktail party problem’ in the sense that the independent signals are like voices of many individuals, while the mixed signals **s**_1_, **s**_2_,… are picked up by microphones in various places. We know the consumers need to listen only to a few of these voices, but we do not know in advance which voices these will be. Each consumer receives its ideal output as a training signal. The loss function for the whole network is 

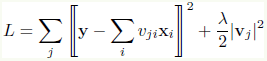

This simulation shows how the neural marketplace framework can be used to tackle such a problem:

- Each producer uses an unsupervised learning rule to perform *independent component analysis* and recover a single source **b**_i_. Because ICA will find a source at random, most producers will represent distractor sources.
- Each consumer uses a delta rule to find weights **v**_*j*_ that minimise its contribution to *L*, namely 

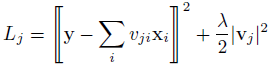
- Each producer *i* infers from its 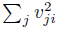 whether the source it recovered is relevant for the consumers. A small 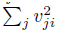 indicates irrelevance, prompting producer *i* to randomise its input weights and seek a different voice to present to the consumers.

**Figure 5:**
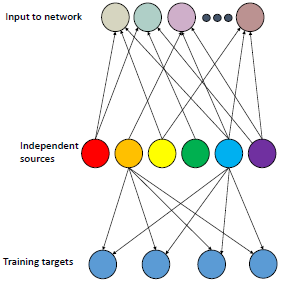
The ‘cocktail party’ problem used to test the PMH algorithm. Inputs to the network (top row) are generated by summing independent non-Gaussian sources (middle row), with dense weights. Training targets (bottom row) are generated as linear combinations of a small subset of the sources.

#### 6.2.1 Problem details

In this simulation, the stimuli are linear combinations of 15 independent signals *b_1_,…, b_15_*. Each *b_i_(t)* is independent and identically distributed, drawn from an exponential distribution. These independent signals are passed through a random mixing matrix **A** before presentation to the network. **A** is orthonormal and each *b_i_* has 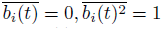. The stimulus vector seen by the producers at time *t* is **s**(*t*) = **Ab**(*t*). Producer *i* has output 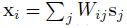.

The target for consumer *j* is 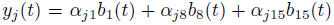, with independently drawn 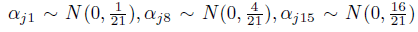. Thus, sources 1, 8, and 15 are useful to the consumers (source 15 very important, source 1 less important), while the other 12 sources are purely distractors.

#### 6.2.2 Implementation details

For the mathematical analyses of section 5, we assumed that the supervised and unsupervised learning rules used by the consumer and producer neurons, ran to completion before the each step of the PHM algorithm. In biology however, this is unlikely to be the case. We therefore test here a slightly different algorithm, in which these gradient-based learning rules occur continuously and simultaneously, and test using simulation how closely this more realistic case conforms to the mathematical theory. This scheme is summarized as algorithm 4:

##### Algorithm 4 Continuous PMH with unsupervised gradient rule for q and supervised gradient rule for v. After a cell proposes a change, it allows max time steps for these two gradient rules to converge before deciding whether to accept or reject the change.

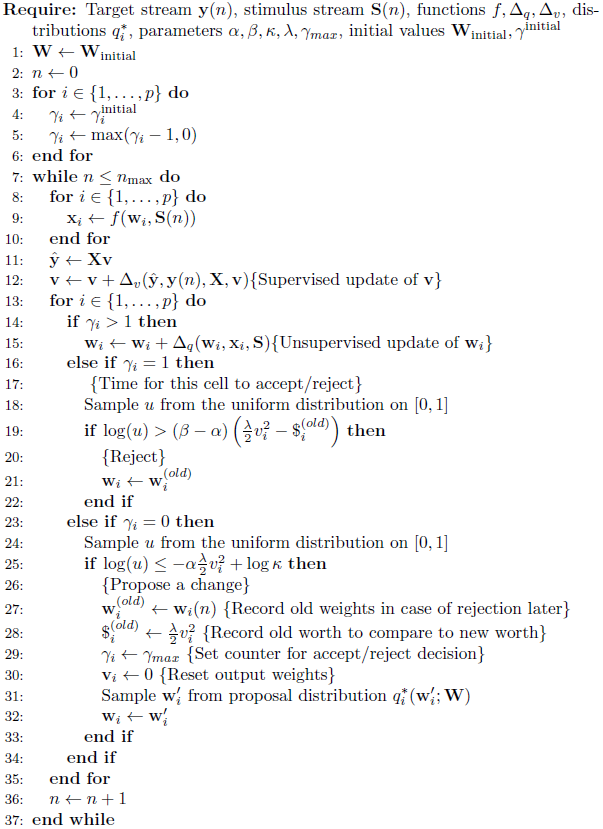

We use algorithm (4) with the following choices of 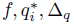, *and* Δ_**v**_:

*f*: the activity of the producers is linear, *x*_*i*_ = **w**_*i*_.**s**.
q_i_^*^: this is uniform on the unit hypersphere 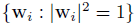.

Δ_q_: producer neurons implement a Hebbian rule to perform ICA. Recall (Hinton and Sejnowski, 1999; Hyvarinen, Karhunen, 2001) that if 
- **b**_1_, **b**_2_ …: are independent and supra-Gaussian, with 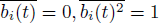,
- *z* is a non-linear, non-quadratic function of 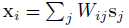, and
- we constrain 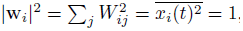,
 then 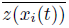 is locally maximised by setting **w**_*i*_ = **a**_*j*_. Thus, unsupervised ICA can be achieved by following the projected gradient of *z* onto 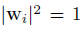. We choose 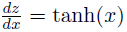and small constant learning rate *μ*, so 

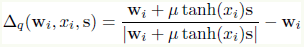

Δ_*v*_: consumer neurons implement a Delta rule to perform ridge regression. Each consumer neuron *j* receives a copy of **y**_*j*_ as a training signal and uses a delta rule to seek appropriate input weights **v**: 

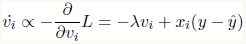

#### 6.2.3 Results

The overall results of the algorithm are illustrated in figure 6. In this simulation, the consumer neuron continuously runs a delta rule. The ICA rule is only enabled after iteration 2000. Initially this leads to a *decrease* in network performance (Fig. 6b), indicating the representation found by Hebbian unsupervised learning alone is actually worse than a random projection. At iteration 4000, the retroaxonal rule is turned on, leading to an immediate improvement in performance. The value the loss function would take if the consumer weights **v** were chosen optimally is very close to the value produced by the delta rule, (Fig. 6b, red points; we compute the optimal **v** analytically from **W** and *α*). The number of cells performing a radical change in input weights spikes immediately after the retroaxonal rule is switched on, and then rapidly decreases (Fig. 6c); note however that the asymptotic value of this is not zero, indicating that a small set of neurons continue to change their representations even after performance has become good. After ICA, neurons tend to represent individual sources, but with no bias toward useful sources; after retroaxonal learning, they are biased to represent useful sources (Figure 6a).

**Figure 6:**
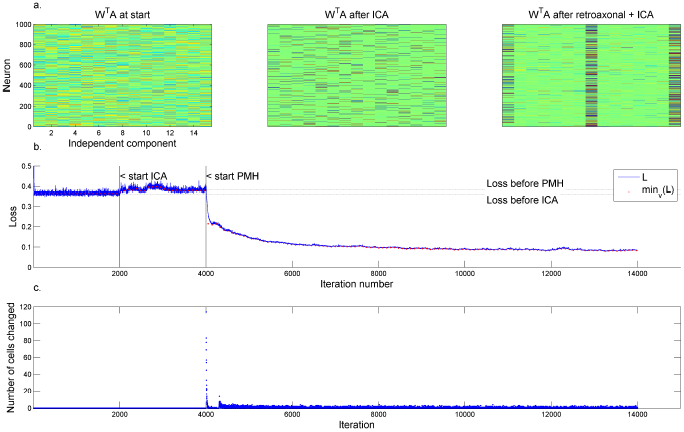
A simulation using PMH for the cocktail party task, and producer neurons learning by unsupervised Hebbian plasticity. *a*. Pseudocolor plot of representations found by the producer cells before learning (left); after ICA alone (middle); and after both ICA and retroaxonal learning (right). *b*. Loss function vs. iteration number (blue curve); red dots indicate the minimum loss function possible for the current producer weights **W**, computed by analytically calculating the optimal **v**. *c*. Number of cells experimenting with large changes to input weights at each iteration.

The worth approximation 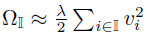 holds with good accuracy in these simulations, for sets of up to 100 neurons, at all stages of the learning rule (Figure 7). For larger sets (500 neurons), the approximations become less accurate. However, even in this case the relationship between predicted worth and actual worth is monotonic. Because the actual consumer weights **v** are learned with a delta rule, they do not take exactly optimal values. Nevertheless, simulations indicate that the worth predicted by these weights closely matches the actual worth value computed by removing the neurons and relearning output weights (Figure 7, blue and red dots), with a poorer fit visible only immediately after the retroaxonal rule has been switched on, reecting the fact that large numbers of cells had recently radically changed input weights. Even at this time however, the relationship between actual and estimated worth is monotonic.

**Figure 7:**
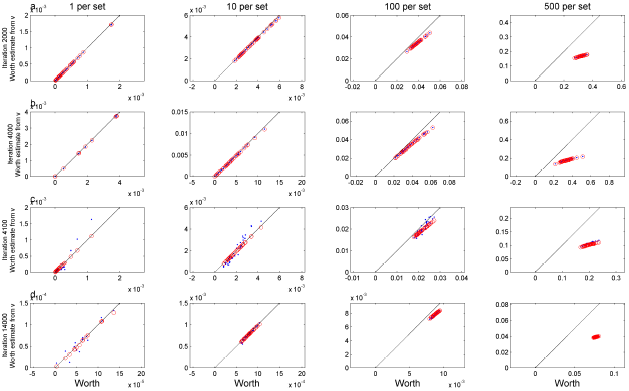
The approximation 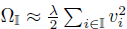 holds with good accuracy during simulations of the PMH algorithm. Each panel shows a scatter plot representation of actual worth of a set of neurons (computed by calculating optimal performance with and without the set) vs. its approximated value. Blue dots indicate the approximation computed from the current weights **v** learned by the delta rule; red circles indicate the values predicted from the optimal **v** derived analytically. Each column of panels shows results for cell sets of different sizes, with every point representing a randomly-drawn set of this many cells. Each row of panels corresponds to a different time point in the simulation: early in running of ICA alone; shortly before the retroaxonal rule is switched on; shortly after the retroaxonal rule is switched on; and substantially after it is switched on.

Increased network performance does not correspond to increased worth of individual cells (Figure 8). After ICA is switched on, the worth of some cells increases. The cells whose worth increases are those whose activity represents important sources such as source 15 (given the highest weights in the consumer training signal). After the retroaxonal rule is switched on, the worth of each individual cell representing source 15 drops dramatically, the network performance improves, and the number of cells representing source 15 increases. The singular value decomposition (section 5.5) explains this evolution of worths: when *p* neurons represent source 15, their output weights will scale as 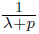 and individual worths scale approximately as 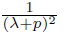. Thus, when large numbers of cells represent source 15, each one’s worth will be lower, while network performance is overall higher. In the economic analogy, this can be likened to the effect of competition on price: when a small ‘oligopoly’ of producer neurons are the only ones manufacturing a product, they are able to charge a high price for it; but competition will drive this price down.

**Figure 8:**
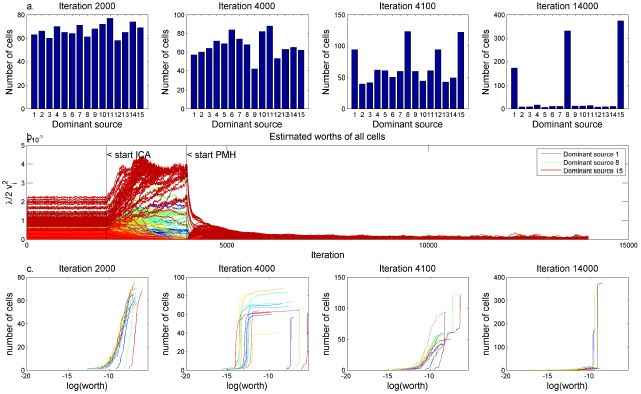
Increased network performance does not correspond to increased worth of individual producer cells. a. Histogram of the number of cells dominated by each source at different iterations. Cell *i* is ‘dominated’ by source *j* if *j* = argmax_*k*_|**w**_*i*_.**a**_*k*_|, where **a**_*j*_ is column *j* of **A**. b. Each curve shows the estimated worth of a single producer cell as a function of iteration number; curves are color coded by dominant sources. c. Unnormalised cumulative distribution of worths, with cells classified by dominant source (color code as in b.). After ICA alone, cells with useful dominant sources have higher worth, but are no more numerous than, those with useless dominant sources. After ICA and PMH (iteration 14000), most cells represent useful sources and their worths are tightly clustered around a common value.

## 7 Discussion

We have outlined a theory for how neurons can self-organize into information processing networks, analogously to how independent actors self-organize in a free-market economy.

The theory rests on the *retroaxonal hypothesis*, which proposes a new form of communication between neurons, that is not included in current computational models of the brain, but is entirely consistent with the known cellular capabilities of neurons. The retroaxonal hypothesis has two components: that *neurons with weak output synapses show unstable input synapses, and that strengthening of a neuron's downstream synapses consolidates recent changes in the same neuron’s inputs*. While direct tests of the hypothesis have not yet been performed, we have outlined in section 2 a powerful case that this process occurs, built from data in the experimental literature.

We described a mathematical formulation of the theory in section 4. This formalism applies to any neuronal network models (e.g. spiking, rate-based, recurrent, feedforward). The formalism rests on the concept of *worth*, defined to be the loss in network performance that would occur if a set of cells fell silent, followed by compensatory plasticity of other synapses in the network. We showed that under certain conditions, a this formalism enables learning equivalent to a parallel form the the Metropolis-Hastings algorithm. The Metropolis-Hastings algorithm is commonly used in engineering applications, as it forms the basis of the popular simulated annealing method for global optimization. Thus, the present scheme represents a that the brain could apply an optimization approach similar to simulated annealing, in a parallel fashion.

It is important to note that the scheme described here is *not* a form of gradient-based optimization. Gradient-based approaches (such as the backprop algorithm) suffer from a problem of local minima, that can be a critical obstacle for large and complex networks. The present scheme — in common with approaches like simulated annealing — avoids this difficulty by including a stochastic component. The network constantly experiments with large, non-gradient changes to individual cells; these changes are kept when they increase a cell’s worth, and may also occasionally be kept when worth decreases. This ability to make non-gradient steps, and to accept occasional decreases in performance, are critical to how simulated annealing avoids becoming stuck in local optima.

The convergence proof presented in section 4 rests on several assumptions, which will in general hold only approximately. One of the key assumptions is additivity of worths: that for a sufficiently small set of neurons, the worth of the set is approximately equal to the summed worths of all neurons in the set. It is this assumption that allows solution to the credit assignment problem. When a large number of neurons make simultaneous changes in their inputs, the ability of individual neurons to compute changes in their worth allows each neuron to identify whether its contribution to network performance improved or worsened. Indeed, assigning a single numerical credit value to an individual neuron is not even possible without an assumption of this form.

We showed analytically that the assumption of approximately additive worths holds, for the case of a simple feedforward linear network (section 5). This analysis also clarified the conditions under which the approximation is accurate. These conditions can be intuitively summarized as a requirement for redundant coding: when large numbers of neurons encode similar features, such that the activity of any one cell can be largely predicted from that of its peers, worth is approximately additive and neurons can estimate their worth by local retroaxonal signaling. Such redundant coding appears to occur universally in cortical and hippocampal networks (e.g (Harris et al., 2003; Luczak et al., 2009; O’Connor et al., 2010; Huber et al., 2012). In the economic analogy, this condition corresponds to a state of competition, as opposed to monopoly. An economy self-organizes for the common good only when the income received by a firm or individual correlates with the benefit of their actions to broader society. A monopoly represents a case of ‘market failure’, where this correlation breaks down, and a firm receives a far greater income than they would if free competition existed. Modern capitalist societies have developed laws to ensure such anti-competitive situations do not occur. We suggest that similar pro-competetive mechanisms may exist in the brain also. In the linear networks we analyzed analytically, we showed that a large weight penalty was sufficient to ensure redundant coding and the accuracy of the approximations.

An important question regards the degree to which the approximations required for the scheme hold in more complex situations. In section 6, we described simulations in which producer neurons used unsupervised learning and consumer neurons used a delta rule, and showed that the approximations required hold extremely well. While feedforward connectivity is a good description of many brain circuits involved in learning (for example the corticostriatal pathway or projections from CA1 to subiculum), it remains to be demonstrated that the required conditions also approximately hold in recurrent networks such as those involved in local neocortical circuits. A mathematical analysis of such recurrent networks will be presented in a forthcoming manuscript.

## Footnotes

[1] The average of the non-zero eigenvalues must be 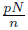 because there are *n* of them and their sum is Trace **X^*T*^X**. To find the interval they lie in, write **X** = **SA** where 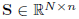 is the matrix of *n* uncorrelated signals and 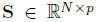 is a random mixing matrix whose entries have mean 0 and variance 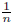. Then 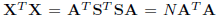. The Marchenko-Pastur law implies that, in the limit where 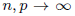 and *n/p* is fixed, the non-zero eigenvalues of *n***A^*T*^A** lie between 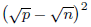 and 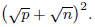 Thus, the non-zero eigenvalues of **X^*T*^X** are in the range 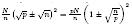.

